# Translationally controlled tumor protein *TCTP* as Peptide Vaccine against *Schistosoma japonicum*: an immunoinformatics approach

**DOI:** 10.1101/466847

**Authors:** Rayan A Abdalrahman, Shima S Ahmed, Mahmoud A Elnaeem, Marwa S Mohammed, Nawraz M Jammie, Israa A Yousif, Wala H Mohamed, Sabreen Y Nasr, Mawadda A Awad-Elkareem, Mohamed A Hassan

## Abstract

Schistosoma japonicum is the most pathogenic causative form of schistosomiasis that causes a major health problem in its endemic countries. Until now, praziquantel is the only drug used to treat Schistosomiasis, but it does not prevent re-infection. So, repetition of the treatment is needed. Moreover, there is no effective vaccine against S. japonicum. Therefore, an urgent need for the development of vaccines is mandatory. This study aimed to analyze an immunogenic protein, Transitionally Controlled Tumor Protein (TCTP) using an immunoinformatics approach to design a universal peptide vaccine against Schistosoma japonicum. A set of 22 of TCTP sequences were retrieved from NCBI database. Conservancy of these sequences was tested and then conserved B cell and T cell epitopes were predicted using different tools available in Immune Epitope Database IEBD. Epitopes having high scores in both B and T cell predicting tools were proposed. An epitope ^129^YEHYI^133^ was predicted as a most promising epitope with good prediction scores in surface accessibility and antigenicity. Besides that, epitopes ^129^YEHYIGESM^137^ and ^92^YLKAIKERL^100^ were predicted as the most promising epitopes concerning their binding to MHC I and MHC II allele respectively. The study revealed that our predicted epitopes could be used to develop an efficacious vaccine against Schistosoma japonicum in the future especially epitope YEHYIGESM as it is shared between MHC I and II alleles and overlapped with the most promising B cell epitope. Both in vitro and in vivo studies is recommended to confirm the efficacy of YEHYIGESM as a peptide vaccine.

## Introduction

*Schistosoma japonicum* is blood flukes that can infect human and livestock causing a chronic disease of considerably importance in public health in endemic countries (1–3). It belongs to the Schistosoma genus of the Schistosomatidae family, which includes mammalian Schistosoma such as *S. jabonicum*, *S. heamatobium*, *S. mansoni, S. intercalatum, and S. mekongi*. This parasite infects human through skin from fresh water infected by larval of parasite that released from snail and lives in the blood vessels causing immune reaction and progressive damage to organs (4–6). Among all species, *Schistosomiasis japonica* is the most pathogenic type since it releases high number of eggs by adult female worm, which deposited in liver, intestine and other organs and induce reaction of parasite with organ (7, 8). Currently, schistosomiasis is endemic in around 74 countries, and more than 200 million people infected by its worldwide (4, 6, 9–12). Schistosomiasis japonica is still endemic in China, Philippines and to some extent Indonesia despite many controlling programs conducted (13, 14).

Till date, praziquantel (PZQ) is the only drug that has been used to treat Schistosomiasis since 1970, but it does not prevent re-infection. So, repetition of the treatment with PZQ is required at frequent intervals (15–19). Additionally, poor activity of praziquantel against immature schistosome, its limited use for patients developed hepatosplenic lesion, and the potential of drug resistance development prohibit influential control depending on chemotherapy alone and necessitate develop and use of vaccines as a key and complementary component beside other integrated approach for the disease control and elimination (4, 9, 15, 19–24).

In previous studies, the use of recombinant proteins such as, Paramyosin (sj97), Thyroid hormone receptor beta (*SjTHRβ*) induced 33–34%, 27% and 33% reductions of the worm burden in vaccinated animals respectively (25–28), while DNA vaccines like *Sj26GST* induced 30% reductions of the worm burden and 45% reduction of the liver egg burden in vaccinated mice (28, 29). Still, no vaccine was able to induce good protection, thus searching for additional vaccine candidates is needed (21, 28, 30). *Schistosoma japonicum TCTP* has been identified as a potential target for vaccine design. According to proteomic analysis, Translationally Controlled Tumor Protein (*TCTP*) has been identified in Schistosomulum adult life cycle stage of *S. japonicum* and *S. mansoni*, the stage where the parasite is found in circulatory system and release eggs. It is a highly conserved protein responsible for immunogenic response inside the host body through its mechanism such as histamine releasing and regulation of B cell reactivity (4, 9). Furthermore, a significant protection against *Schistosoma japonicum* was obtained after inoculating mice with TCTP and adjuvant (CpG) (31). So, *Schistosoma japonicum TCTP* was considered a good target for vaccination. Therefore, the aim of this study is to predict effective B and T cell epitopes from *S. japonicum TCTP* protein by means of an immunoinformatics approach.

Recently, immunoinformatics approach has been accepted as a universal tool in the field of vaccine development. This method can help in the prediction of appropriate epitopes for designing an efficacious epitope-based peptide vaccine, and offers high degree of confidence for the prediction of epitopes, as an epitope selection is a critical step in the design of an epitope-based peptide vaccine (32–37). Our present study is the first study using in silico approach to predict all possible B and T cell epitopes from *Schistosoma japonicum TCTP* protein.

## Materials and Methods

### An outline of the methodology used in this study has been represented in Figure 1

**Figure 1.**
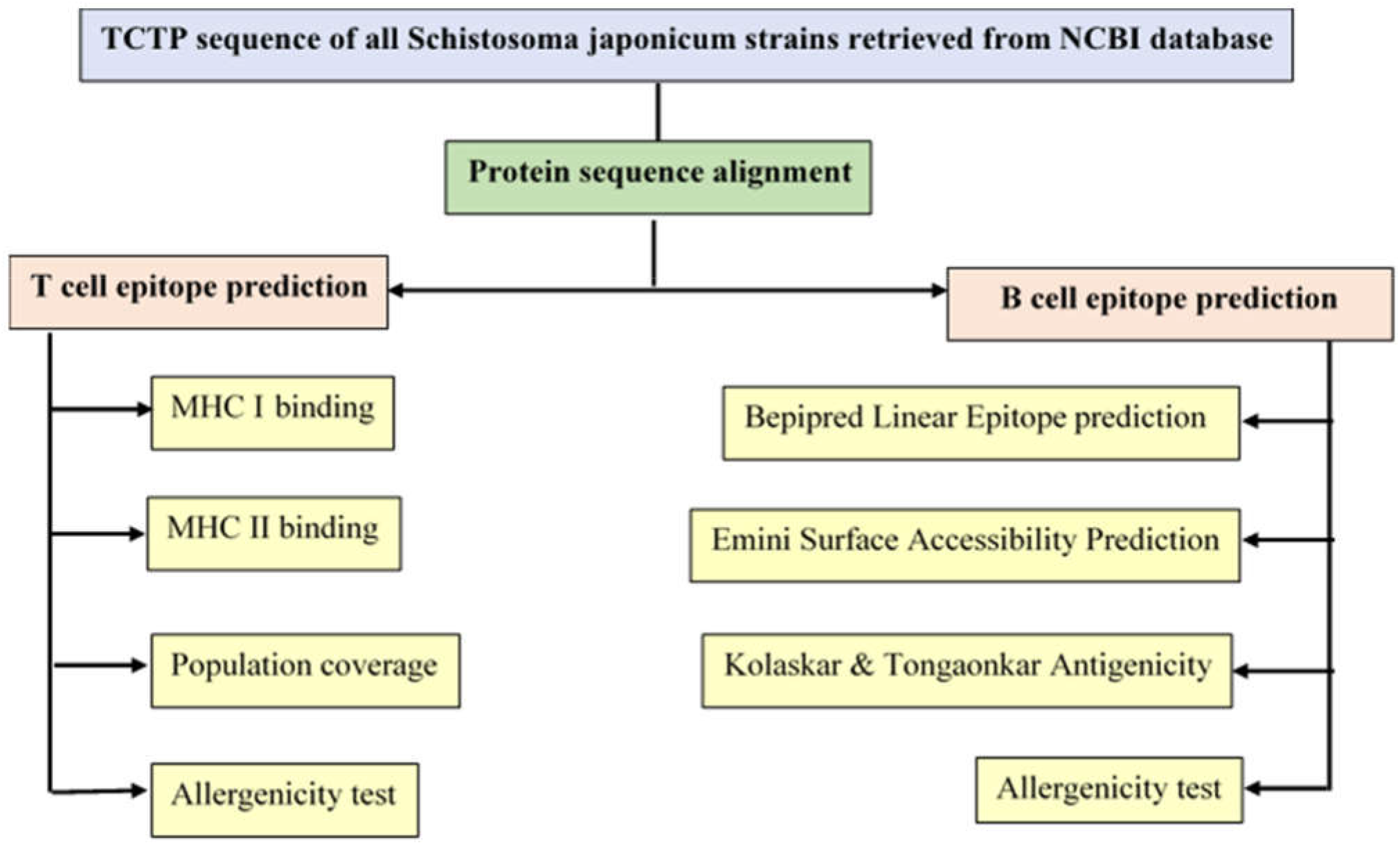
Flowchart representing an immunoinformatics approach used in the prediction of potential B and T cell epitopes in order to develop peptide vaccine against *Schistosoma japonicum*.

### Protein sequence retrieval

A series of 22 sequences of *Schistosoma japonicum* translationally-controlled tumor protein (*TCTP*) were retrieved from the National Center for Biotechnology Information database (NCBI) (http://www.ncbi.nlm.nih.gov/protein/) in 9^th^ of December 2017 for immunoinformatic analysis. The retrieved protein sequences with a length of 169 aa were collected from China; retrieved strains and their accession numbers besides collection area are listed in **Table 1**. Protein sequence (**Accession NO CAX83040.1**) was excluded during retrieval from NCBI database, as it has a length of 196 aa instead of 169 aa.

**Table 1.**
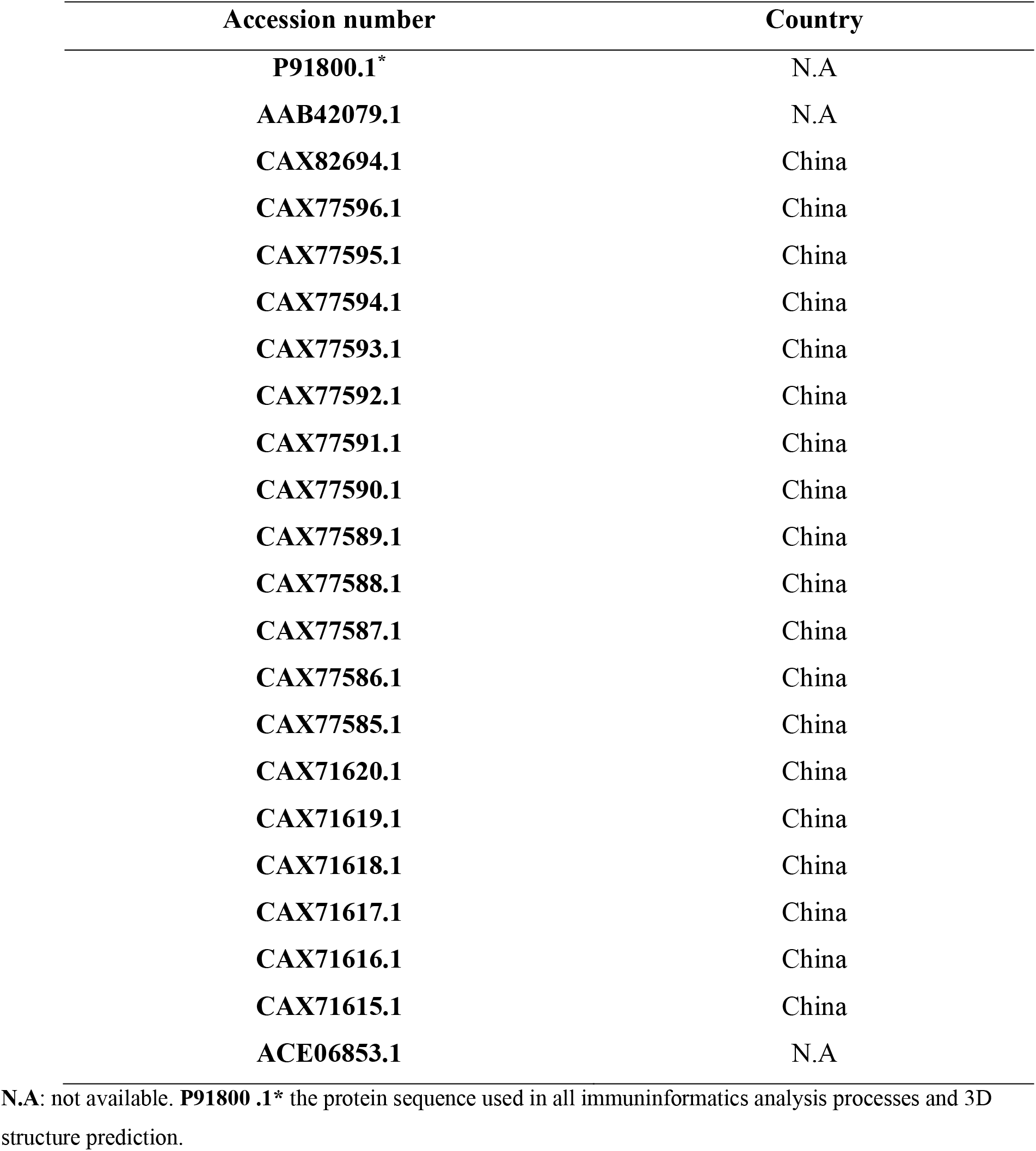
Parasite Strains retrieved with their Accession numbers and area of collection.

### Phylogenetic analysis of the protein sequences

The retrieved sequences were subjected to phylogenetic analysis so as to determine the origin of each strain using Phylogeny.fr software (http://www.phylogeny.fr) (38).

### Protein sequences conservancy analysis

The retrieved sequences were aligned using BioEdit sequence alignment editor (v7.2.5) to get the conserved regions with ClustalW as implemented in the BioEdit program (39). Only conserved regions of the protein sequences were selected for immunoinformatics analysis.

### B cell epitopes prediction

As a result of interaction between the B-cell epitopes and the B-lymphocyte, the latter is differentiated into antibody-secreting plasma cell and memory cells. The main common feature of B-cell epitopes is a surface accessibility and antigenicity (40, 41). Therefore, B cell epitopes were analyzed using different prediction tools from Immune Epitope Database IEDB analysis resource (http://www.iedb.org/bcell/) (42). Bepipred linear epitope prediction tool (43) was used to predict the linear B-cell epitopes from with a threshold value of 0.064. The conservancy of predicted epitopes were tested using BioEdit program and only conserved epitopes were selected for further analysis. After that, Emini surface accessibility prediction tool (44) and Kolaskar and Tongaonkar antigenicity method (45) were used to predict surface accessibility and antigenicity with threshold values of 1.000 and 1.017 respectively. Epitopes which pass these tests were predicted as B cell epitope.

### T cell epitopes prediction

T-cell epitopes are short, linear peptide sequences that are recognized by T cell receptor through their binding with major histocompatibility complex (MHC) to initiate the T cell response (46–48).

### Binding predictions for MHC class I

IEDB MHC I binding prediction tool (http://tools.iedb.org/mhci/) was used to predict the peptide binding to MHC1 molecules. The artificial neural network (ANN) method was used to predict the attachment of cleaved peptides to MHC molecules (42, 49). All epitope lengths were set as 9 mers before prediction. Conserved epitopes that bind to MHC I alleles at score equal or less than 500 half-maximal inhibitory concentrations (IC_50_) were selected for further analysis (50, 51).

### Binding predictions for MHC class II

To predict binding of peptides to MHC II, IEDB MHC II binding prediction tool (http://tools.iedb.org/mhcii/) was used. Human allele references set were used to predict the binding (52, 53). As MHC II groove can bind peptides having different lengths, epitopes binding prediction is more challenging and less accurate than MHC I binding (54). Similarly to MHC I binding analysis, NN-align method was also used to analyze MHC II binding core epitopes (55). Conserved epitopes that bind to alleles at score equal or less than 1000 half-maximal inhibitory concentration (IC_50_) were selected for further analysis (50).

### Population coverage calculation

The Population Coverage tool provides a perception about the efficacy of epitopes to regional and global populations (56). Human population coverage for all predicted MHC I and MHC II epitopes was tested against the whole world population, northeast Asia, and southeast Asia - as places where *S. japonicum* is endemic- with the selected MHC I and MHC II interacted alleles using IEDB analysis resource for population coverage calculation (http://tools.iedb.org/tools/population/iedb_input) (57).

### Assessment of epitope allergenicity

AllerTop v.2.0 (http://www.ddg-pharmfac.net/AllerTOP/) was used to predict the allergenicity of candidate B cell, MHC I and MHC II epitopes. The candidate epitopes were analyzed as either “probable allergen” or “probable non-allergen” meaning that the epitope either cause or does not cause specific IgE production and hypersensitivity responses respectively (58).

### Homology modeling

3D structure of the protein sequence of *Schistosoma japonicum TCTP* was sent to Raptor X (http/www.raptor.uchicago.edu) to predict 3D structure of the protein and then treated with UCSF Chimera (version 1.10.2) to visualize the 3D structure. Furthermore, predicted B cell epitopes as well as all predicted T cell epitopes were verified in the structural level (59, 60).

## Results

### Phylogenetic analysis of the protein sequences

The relationship between retrieved strains of *TCTP* of *Schistosoma japonicum* is shown in **Figure 2**. The phylogenetic tree divided into two groups. One group contained all strains except strain **CAX71615.1** which found in another group. Therefore, protein strain **CAX71615.1** was outgroup and the rest of strains have a common ancestor. Moreover, strains **AAB42079.1** and **P91800.1** were last strains result from an evolution process.

**Figure 2.**
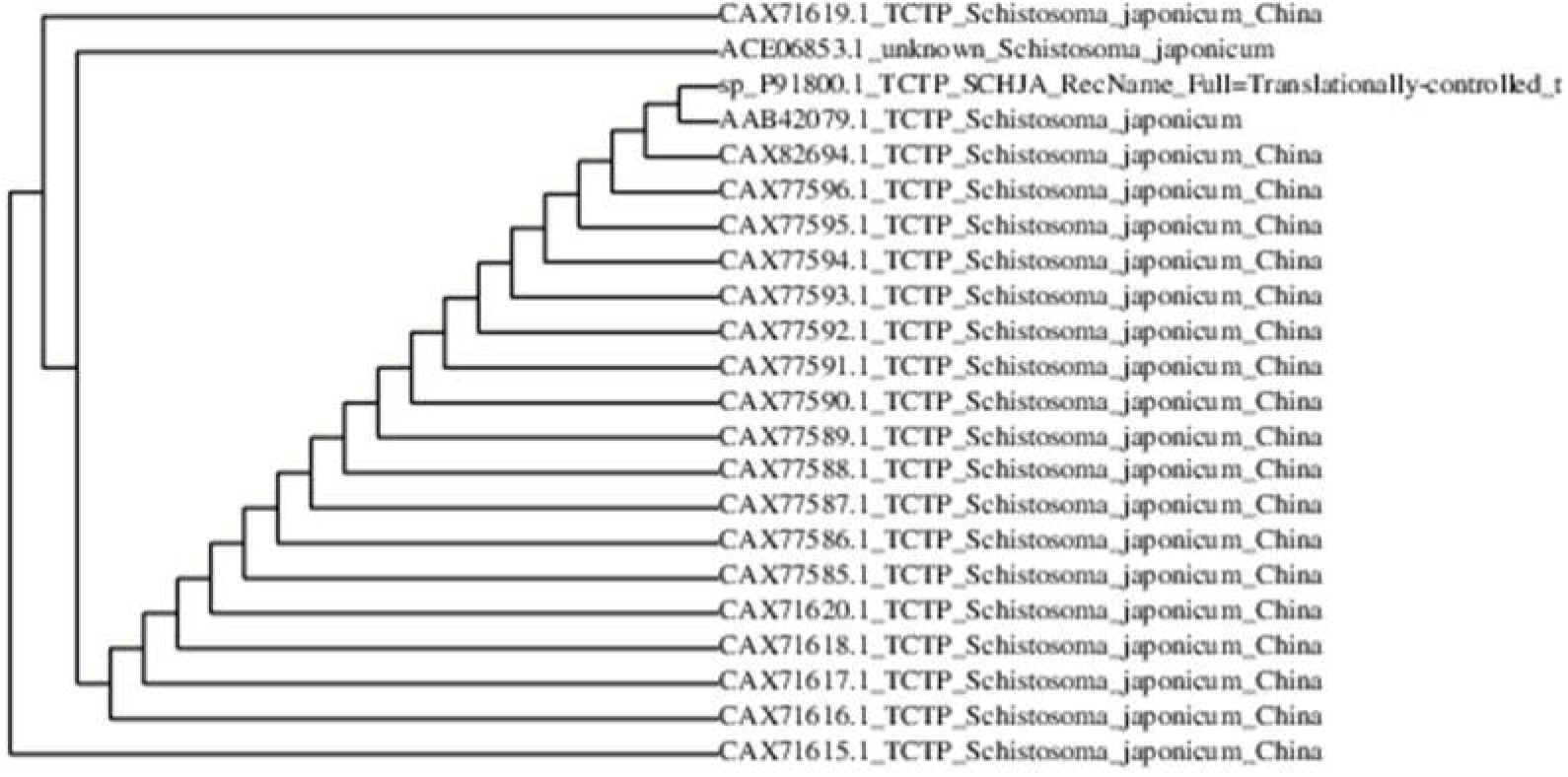
Phylogenetic tree of the retrieved sequences of TCTP of *Schistosoma japonicum* using Phylogeny.fr software.

### Protein sequences conservancy analysis

The use of conserved regions would be efficient in providing broader protection across multiple strains, than epitopes derived from highly variable regions (61, 62). Therefore, retrieved sequences were subjected to conservancy analysis. All the retrieved sequences are highly conserved except two strains **CAX71619.1** and **CAX71615.1** which have mutation at position 127 and 36 respectively as shown in **Figure 3**.

**Figure 3.**
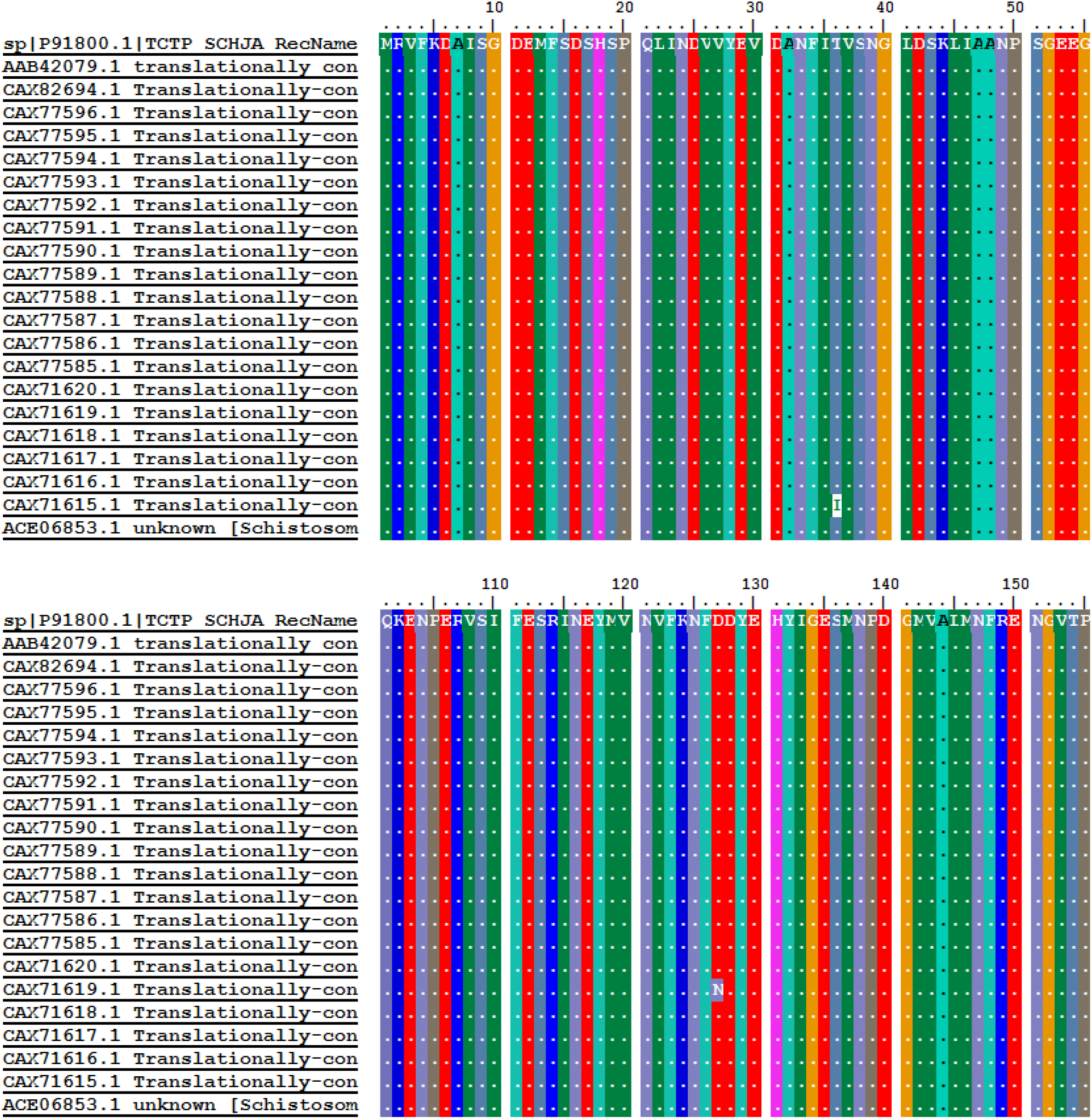
Multiple sequence alignment of whole length amino acid of 22 TCTP strains using BioEdit program (v7.2.5); The highlighted regions represent the mutated region; Dots represent the conservancy between amino acid sequences.

### B cell epitopes prediction

The protein sequence was analyzed using Bepipred linear epitope prediction, Emini surface accessibility and Kolaskar and Tongaonkar antigenicity methods in IEDB to predict the linear epitope, surface accessibility and to assess the antigenicity respectively. Any value equal or greater than the default threshold 0.064 for linearity, 1.000 for surface accessibility and 1.017 for antigenicity were considered as B cell epitope, **Figure 4–6**. Conserved predicted B cell epitopes are listed in **Table 2**. Three linear peptides **DYEHYI**, **DYEHY**, and **YEHYI**, who have higher prediction scores in surface accessibility and antigenicity, were selected as candidate epitopes, their position in structural level is presented in **Figure 7.**

**Table 2.**
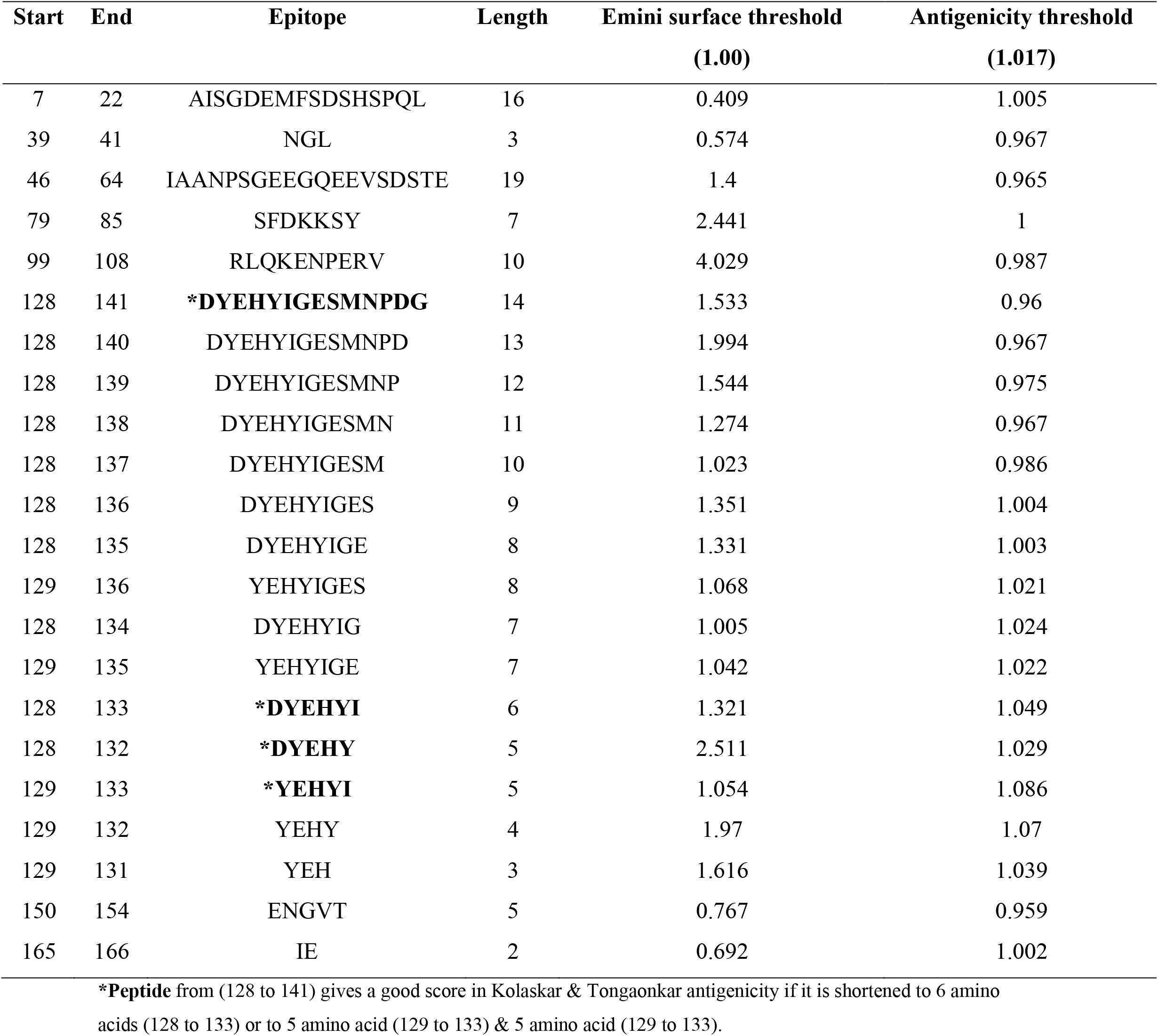
list of B cell epitopes predicted using different scales of *Schistosoma japonicum*.

**Figure 4.**
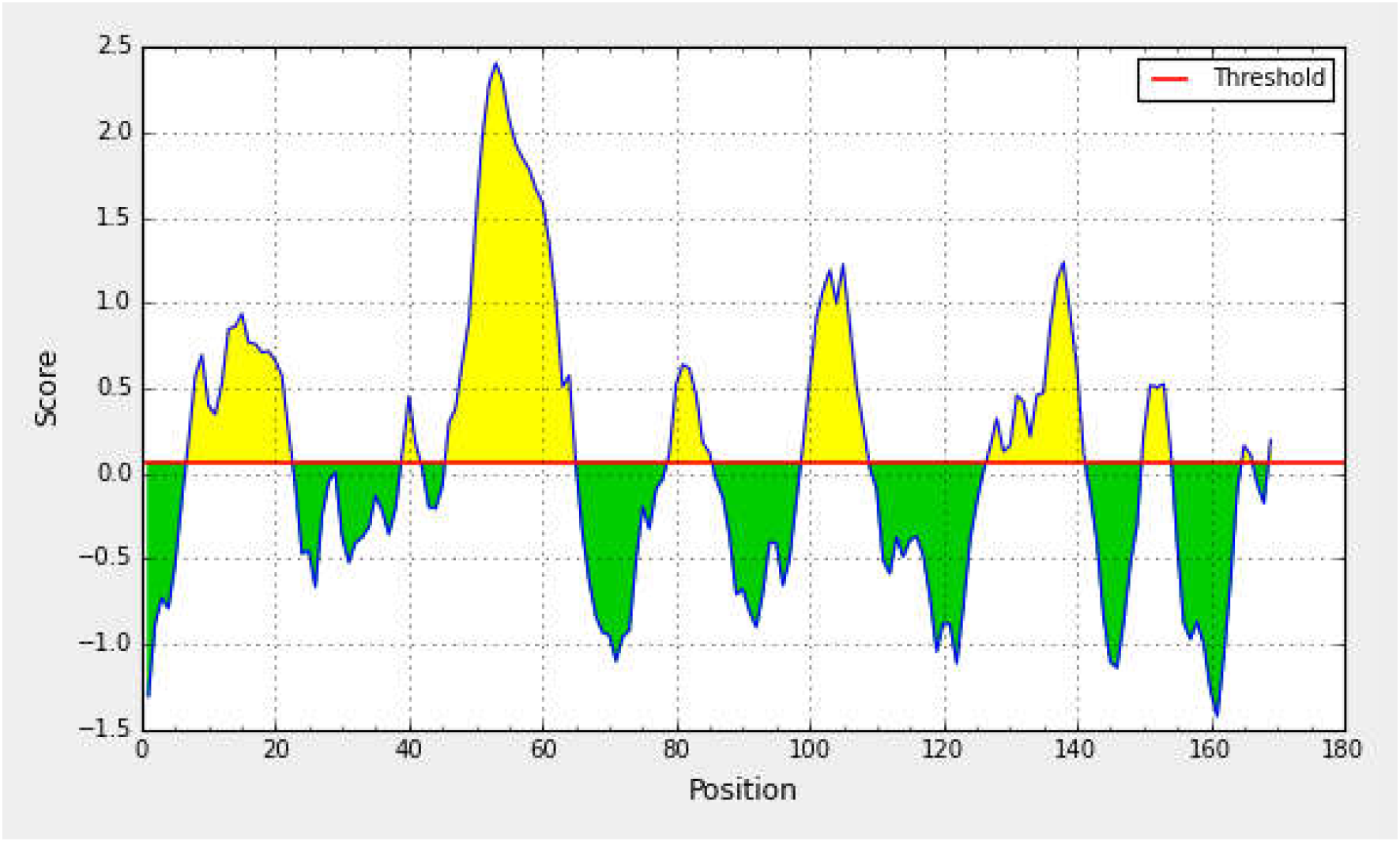
Results of IEDB Bepipred Linear Epitope Prediction tool (**Average**: 0.064 **Minimum**: − 1.425 **Maximum**: 2.407). Yellow areas above threshold (red line) are proposed to be a part of B cell epitopes and the green areas are not.

**Figure 5.**
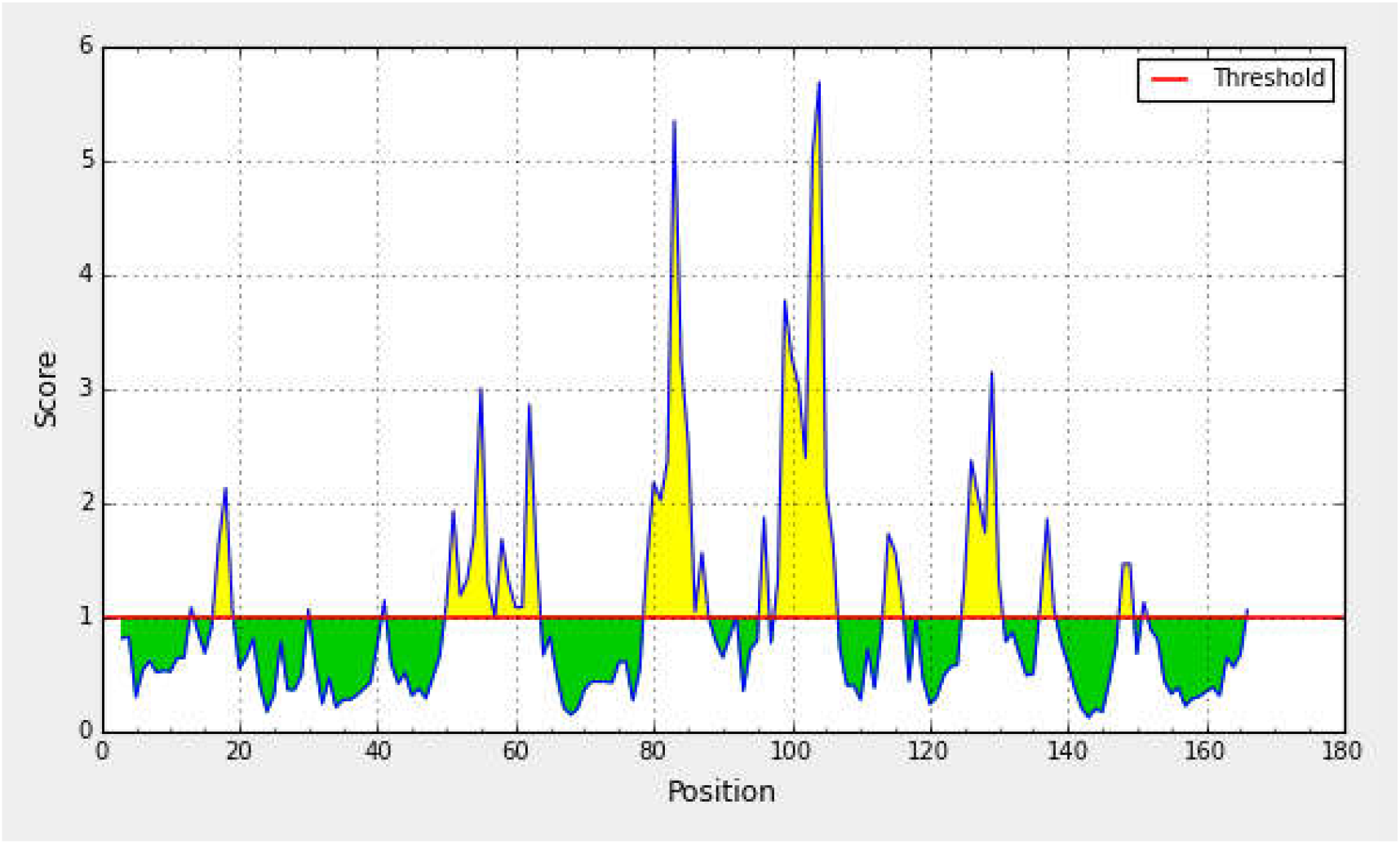
Results of IEDB Emini Surface Accessibility Prediction tool (**Average**: 0.064 **Minimum**: − 1.425 **Maximum**: 2.407). Yellow areas above threshold (red line) are proposed to be a part of B cell epitopes and the green areas are not.

**Figure 6.**
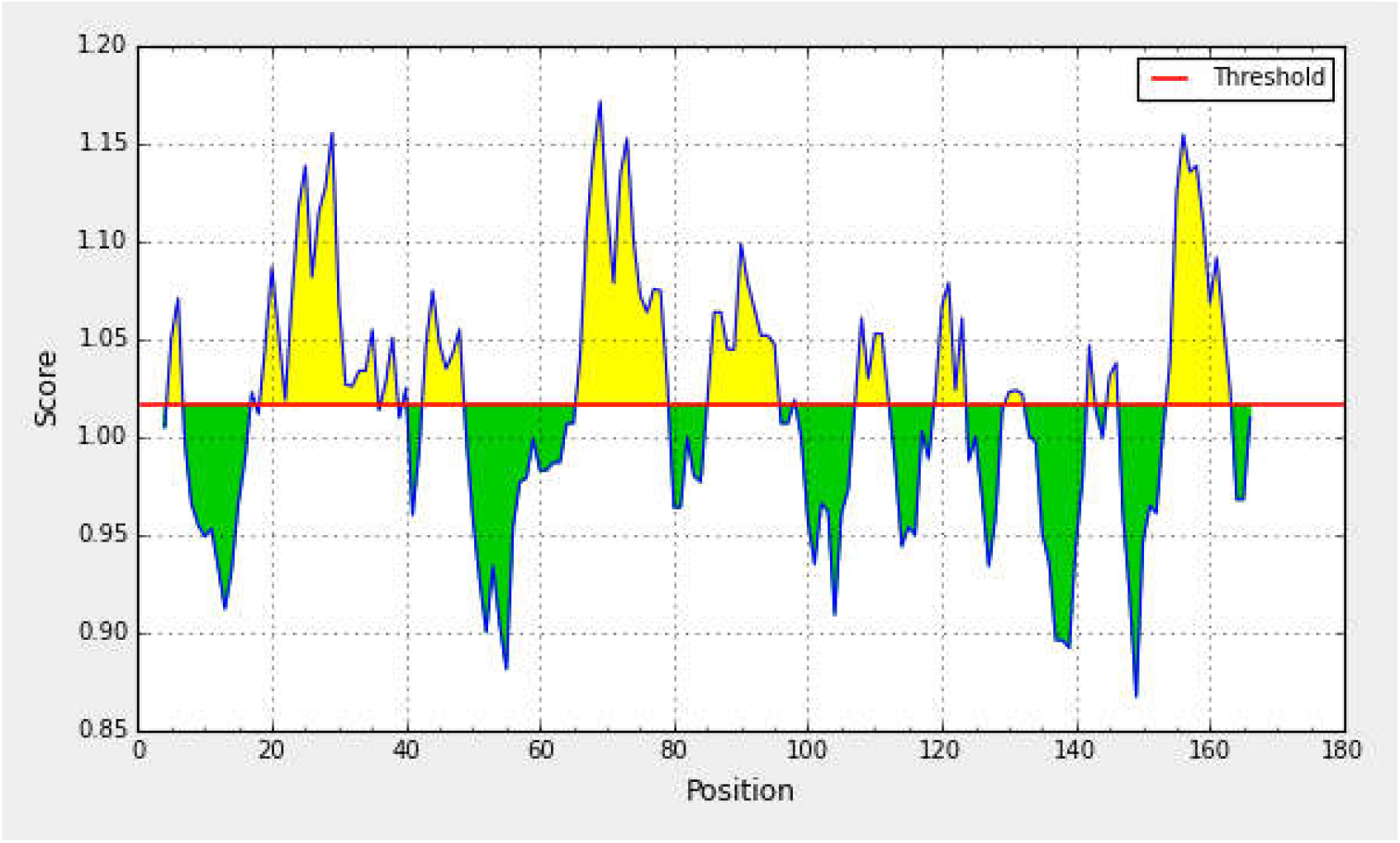
Results of IEDB Kolaskar & Tongaonkar Antigenicity prediction tool (**Average**: 1.017 **Minimum**: 0.867 **Maximum**: 1.172). Yellow areas above threshold (red line) are proposed to be a part of B cell epitopes and the green areas are not.

**Figure 7.**
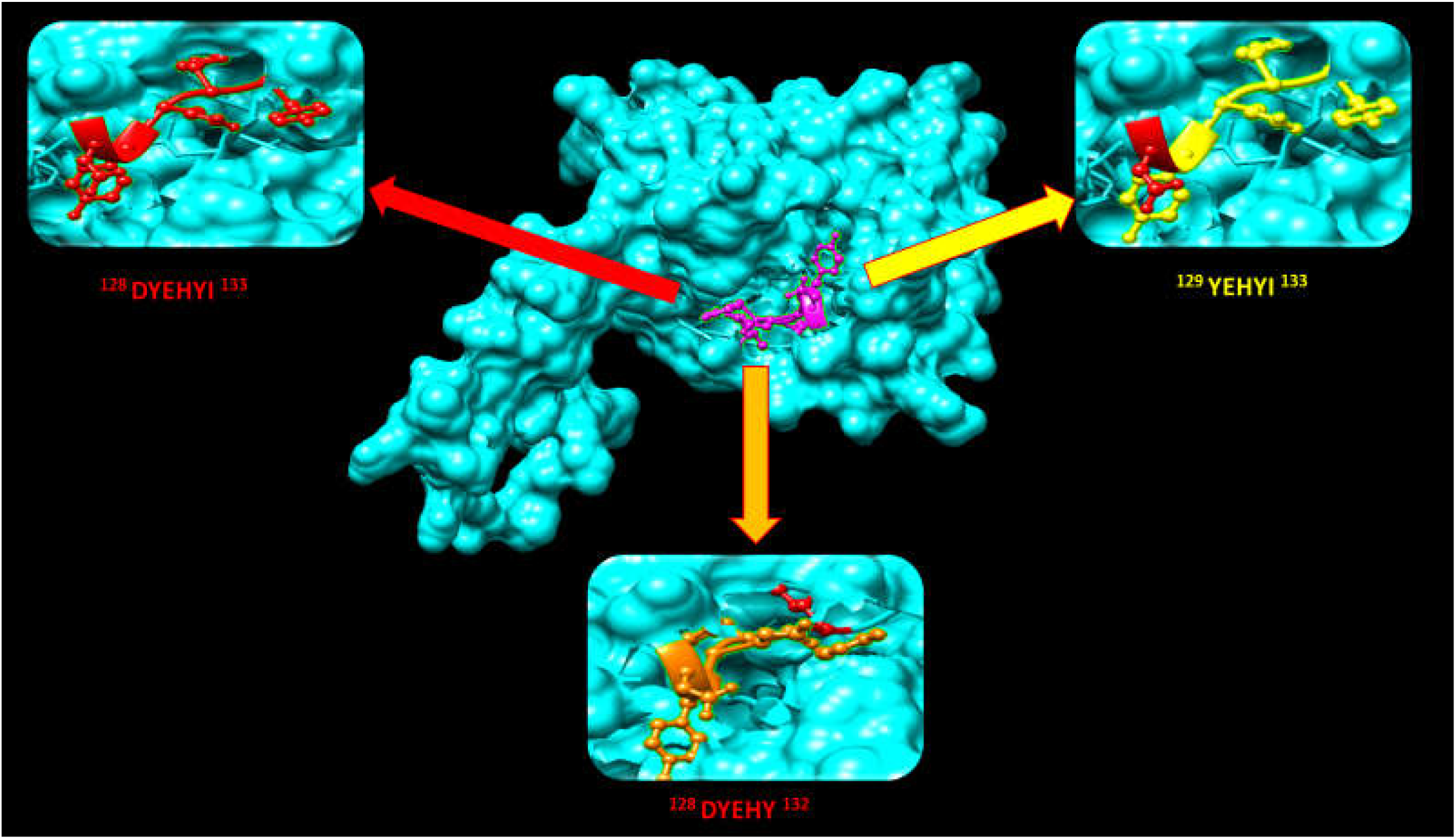
Structural visualization of proposed B cell epitopes of *S. japonicum TCTP* and their overlapping position in the protein from 128 to 133 using UCSF Chimera (version 1.10.2).

### Binding predictions for MHC class I

*Schistosoma japonicum* TCTP was analyzed using IEDB MHC I binding prediction tool. Based on ANN-method, 54 conserved peptides were predicted to interact with different types of MHC I alleles with IC_50_ ≤ 500nM. Four epitopes, **YMVNVFKNF** (118-126), **VVYEVDANF** (26-34), **YEHYIGESM** (129-137), and **FRENGVTPY** (148-156) were found to interact with large number of MHC I alleles with high and intermediate affinity, **Table 3.** Their position in structural level is presented in **Figure 8.**

**Table 3.**
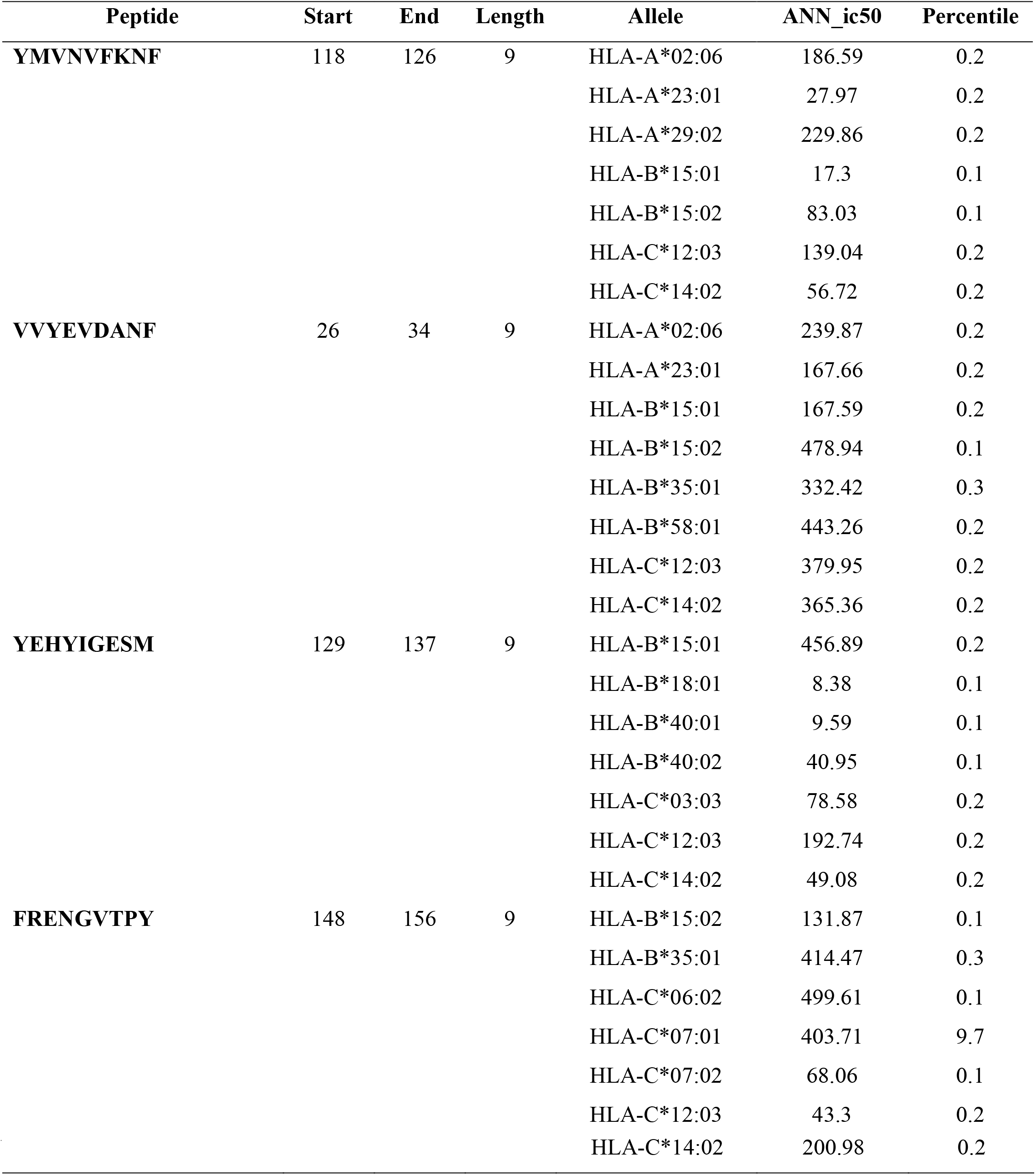
List of epitopes that bind to MHC Class I alleles with high and intermediate affinity

**Figure 8.**
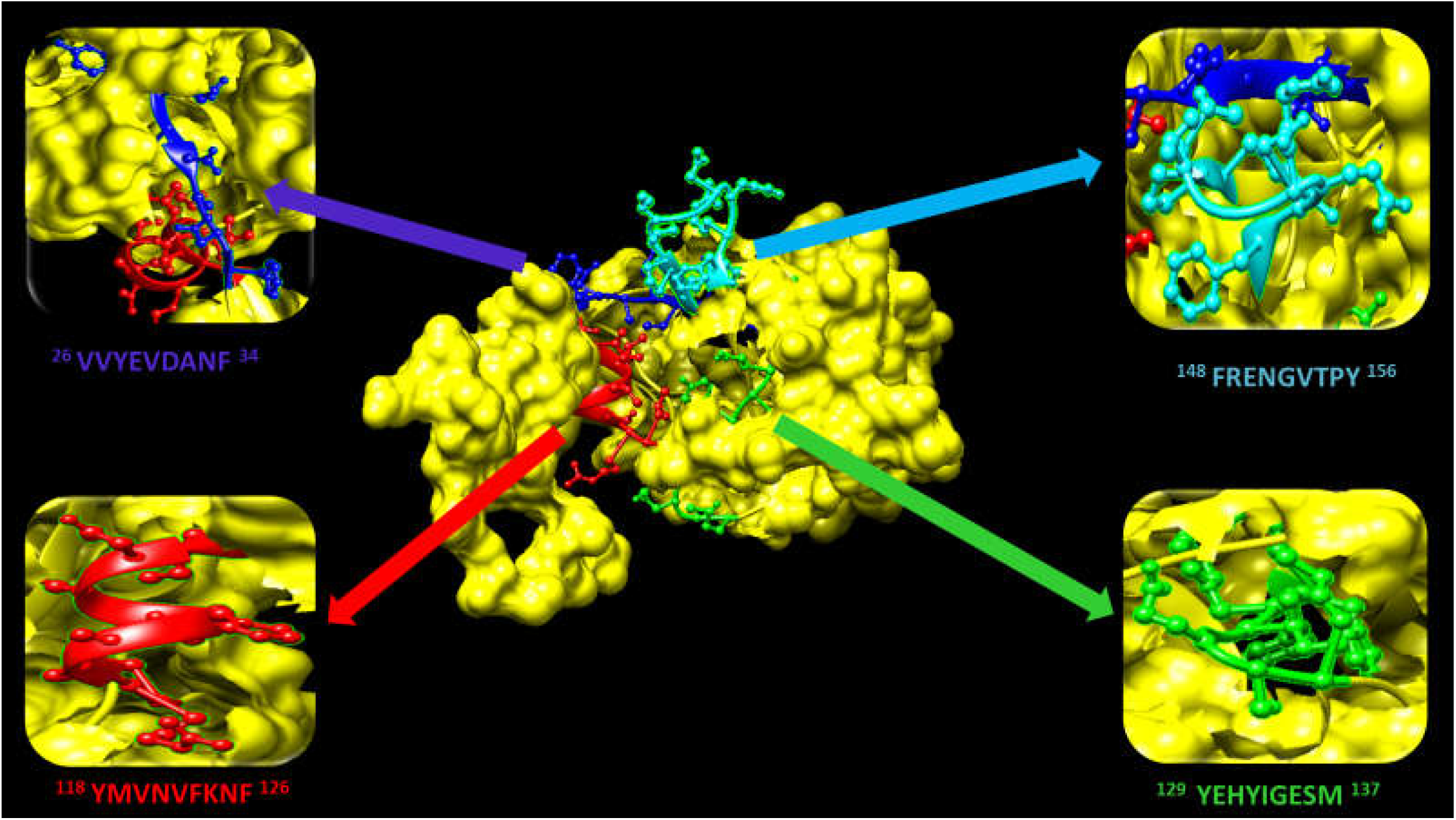
Position of proposed T cell epitopes of *S. japonicum TCTP* that interact with MHC I in structural level using UCSF Chimera (version 1.10.2).

### Binding predictions for MHC class II

96 conserved peptides were predicted to interact with different alleles of MHC II after analyzing the protein sequence using IEDB MHC II binding prediction tool based on NN-align method with an IC_50_ ≤ 1000nM. The core peptides, **IDLVHASRL** had high affinity to interact with twelve alleles, **YLKAIKERL** interacted with sixteen alleles, however **YLKGYLKAI** was found to interact with eleven alleles, **Table 4** and **Figure 9**. Several overlapping between MHC class I and MHC Class II epitopes were found. These overlapping between MHC I and MHC II epitopes are illustrated in **Table 5**.

**Table4.**
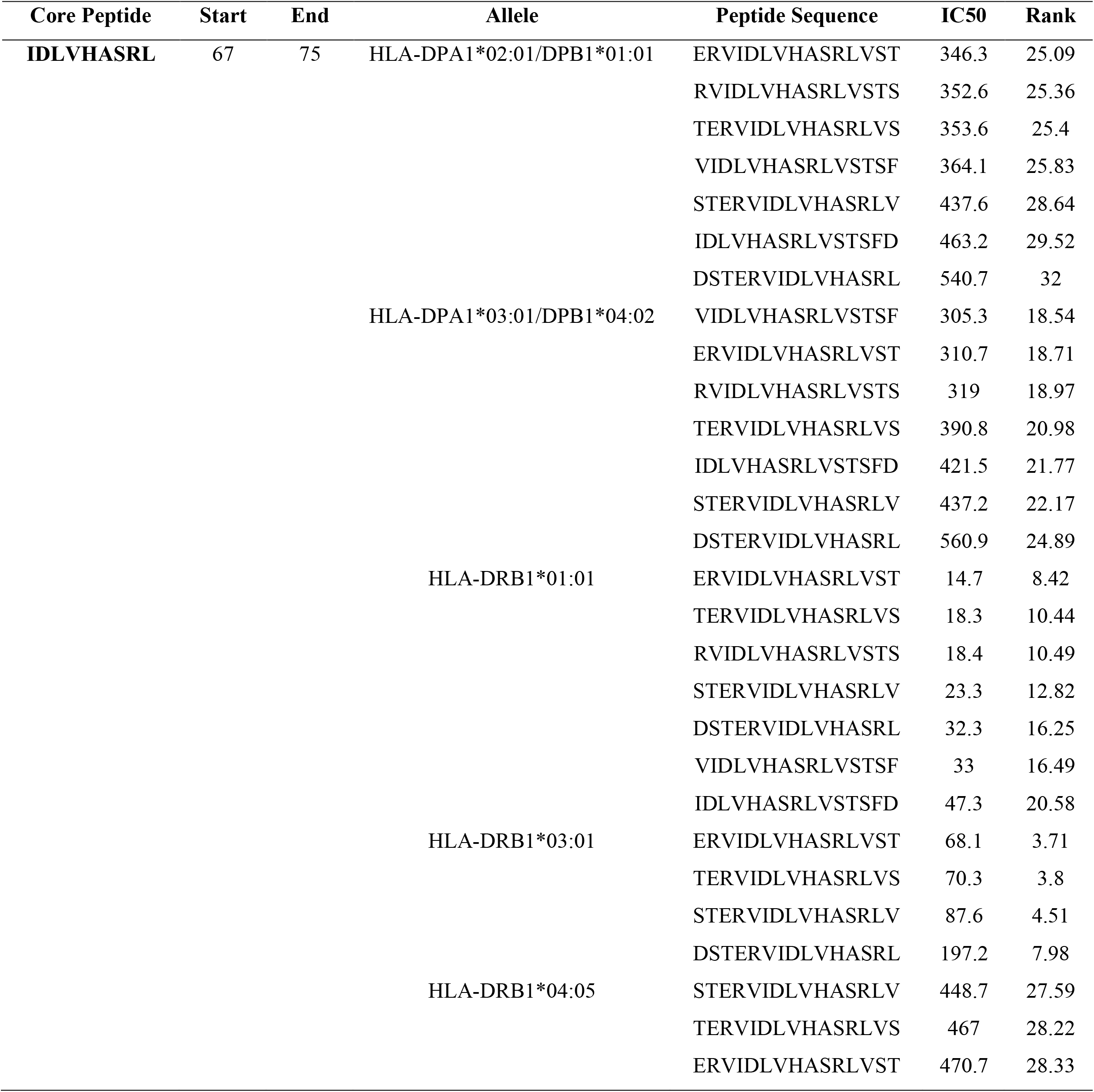
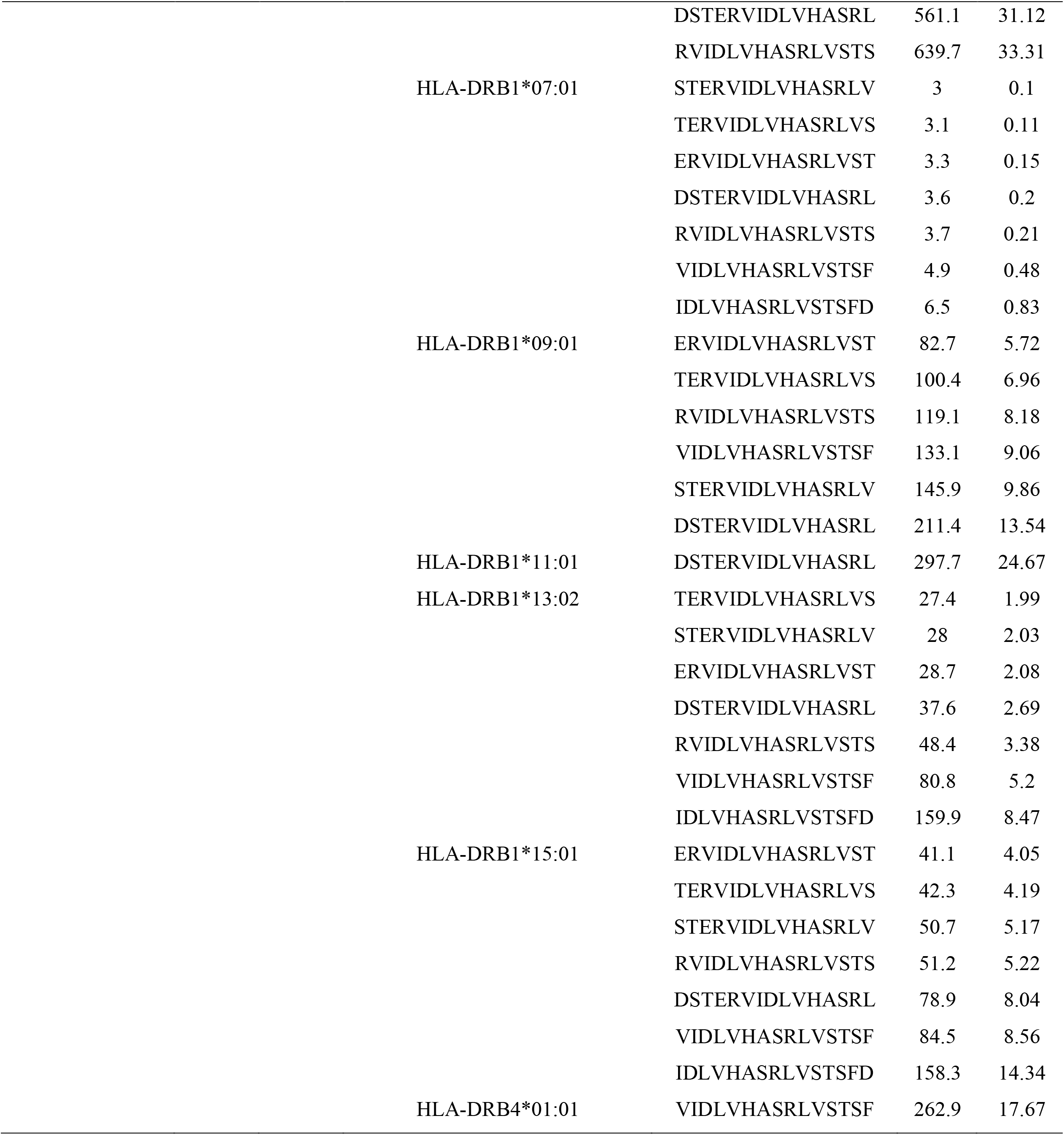
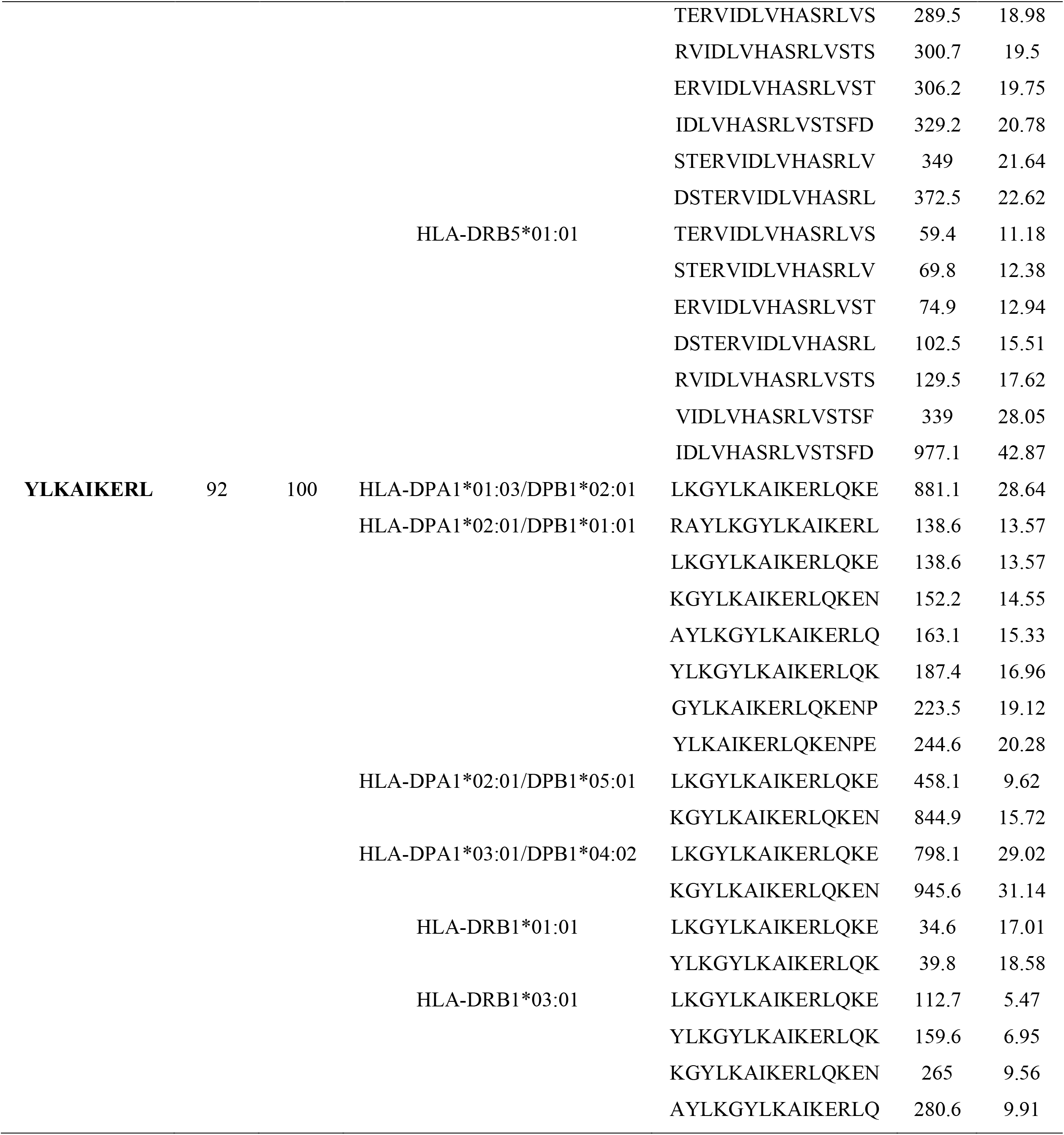
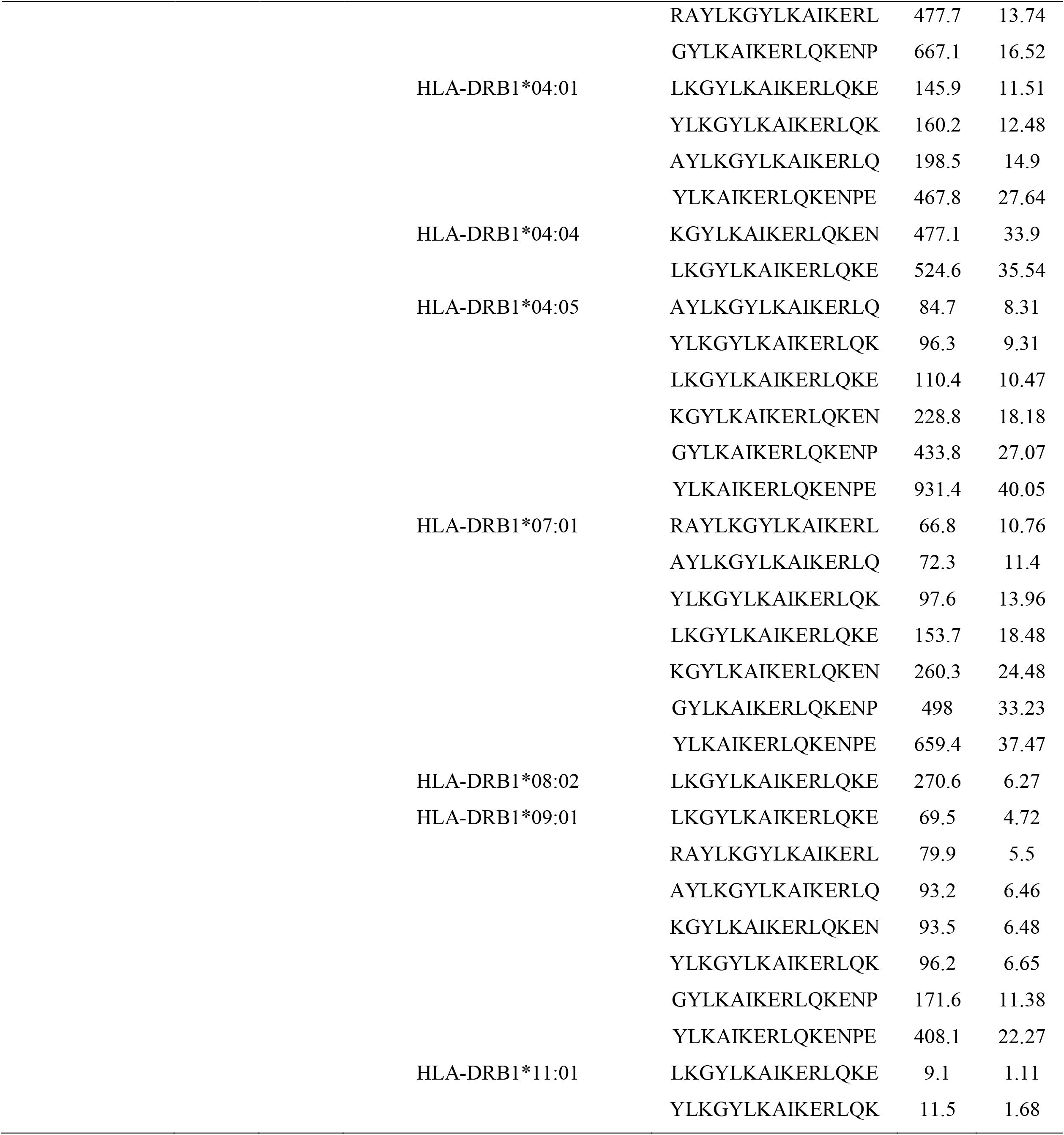
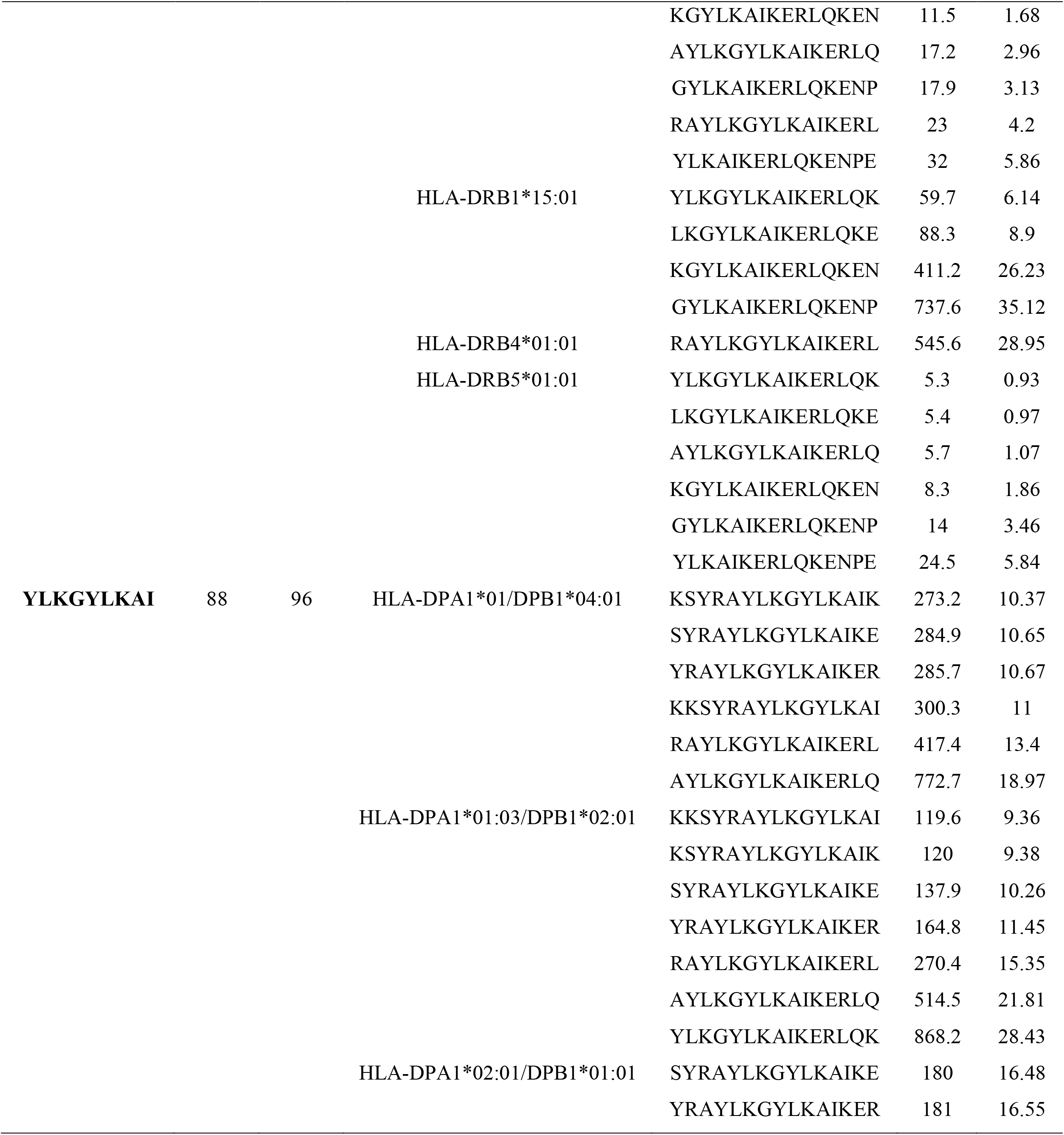
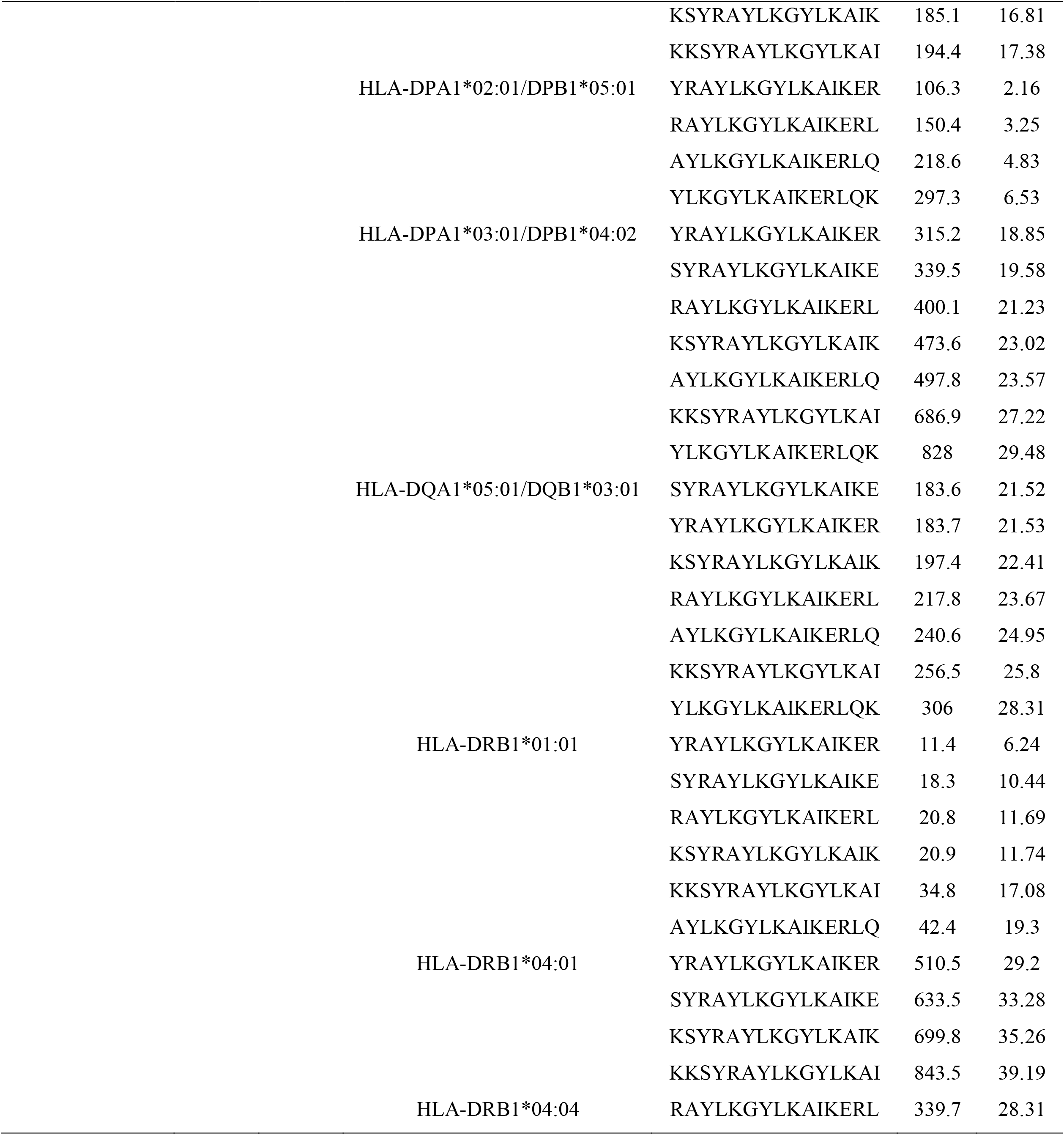
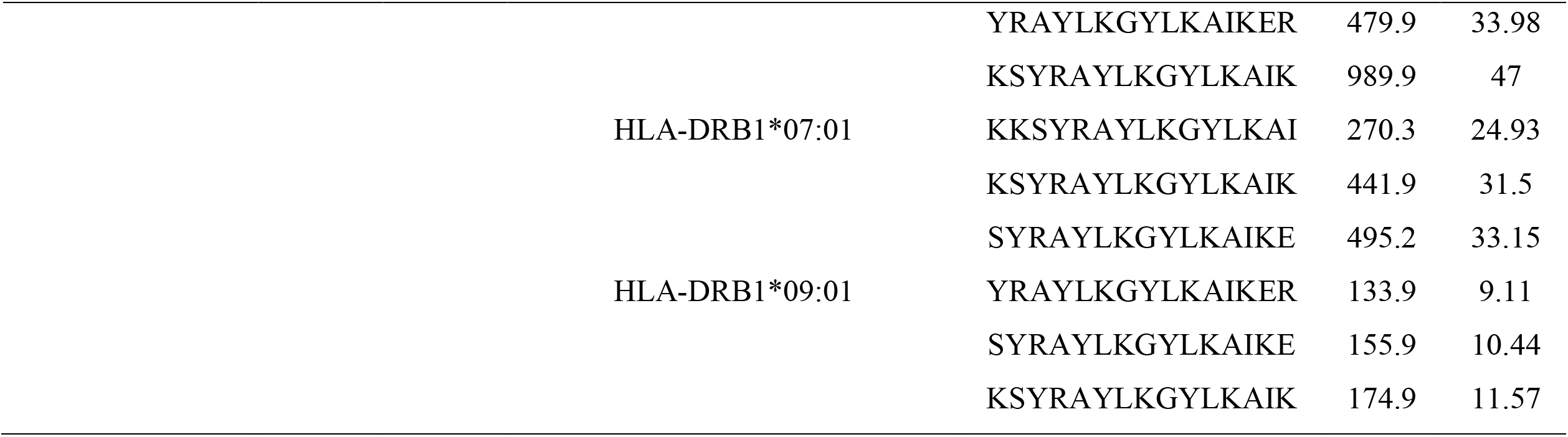
List of epitopes that bind to large number of MHC Class II alleles.

**Table 5.**
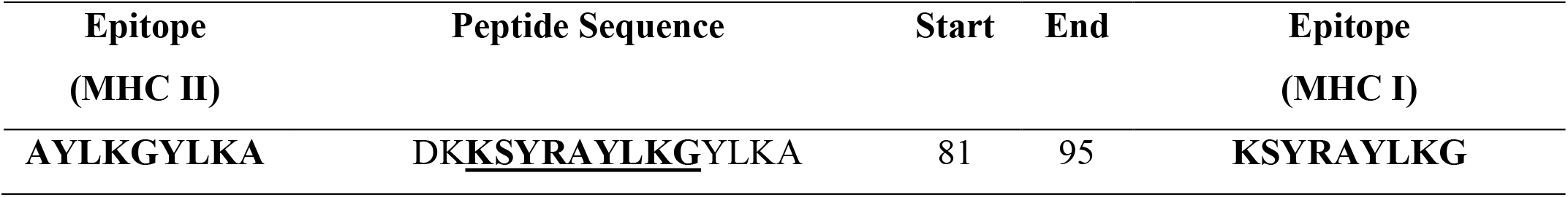
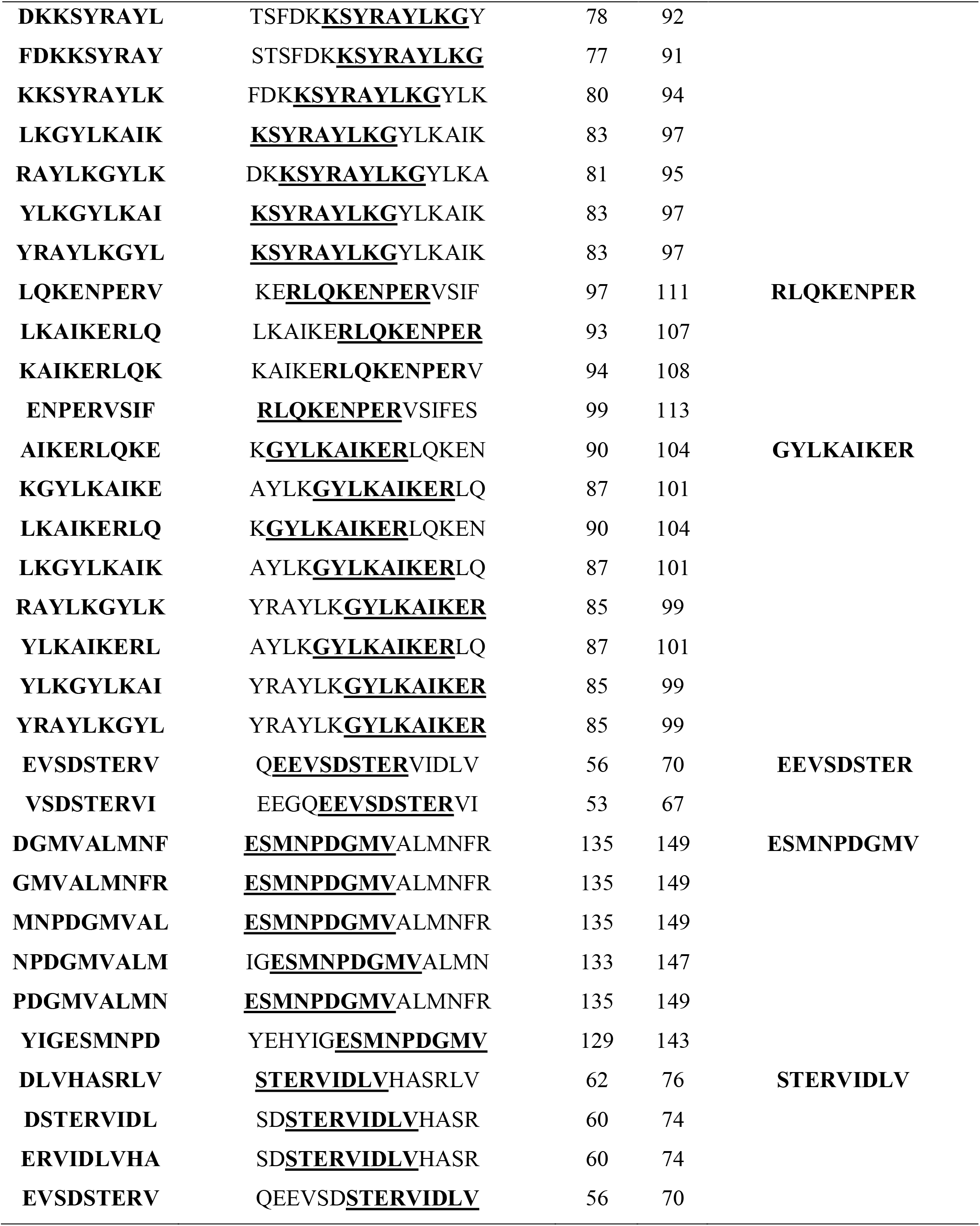
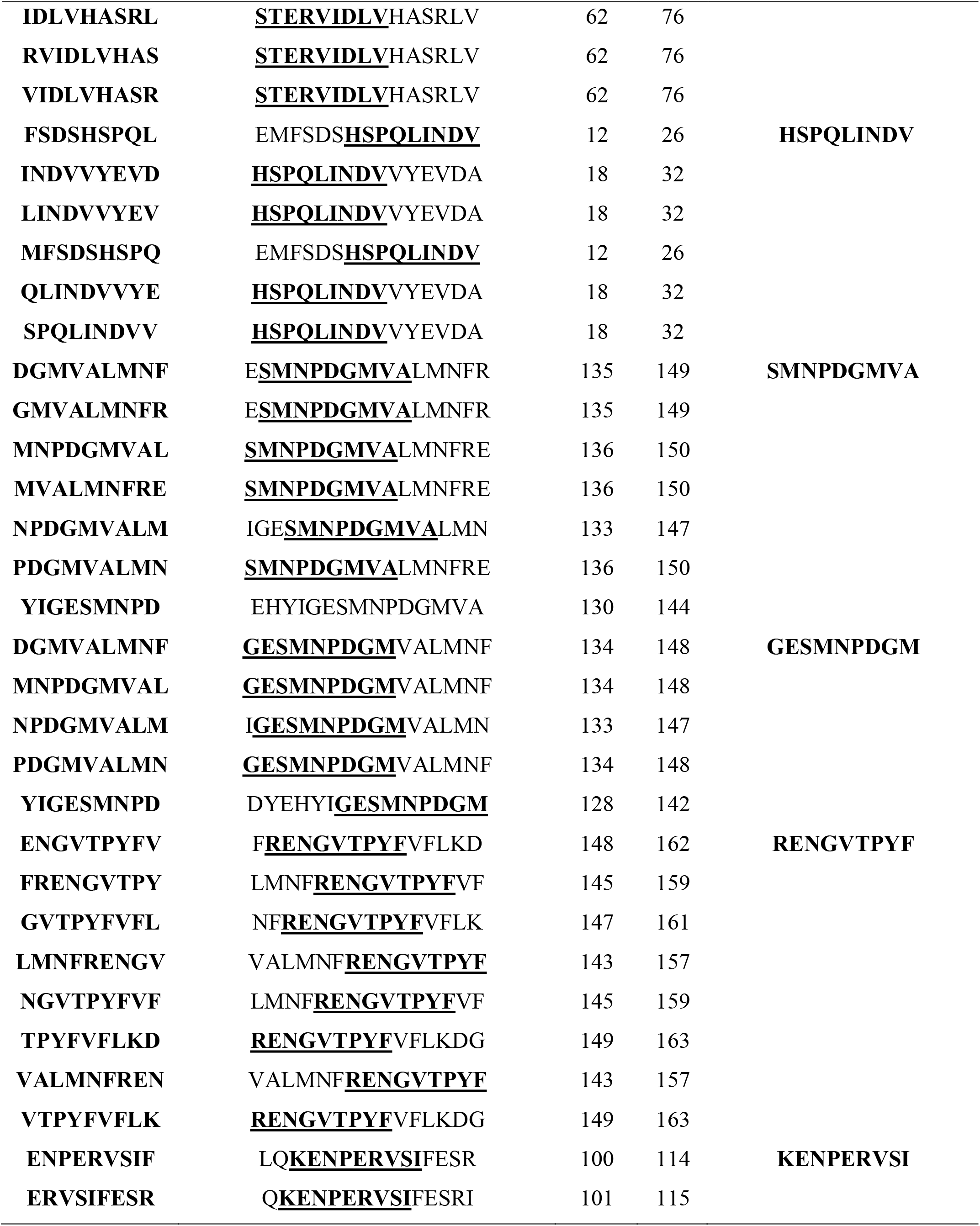
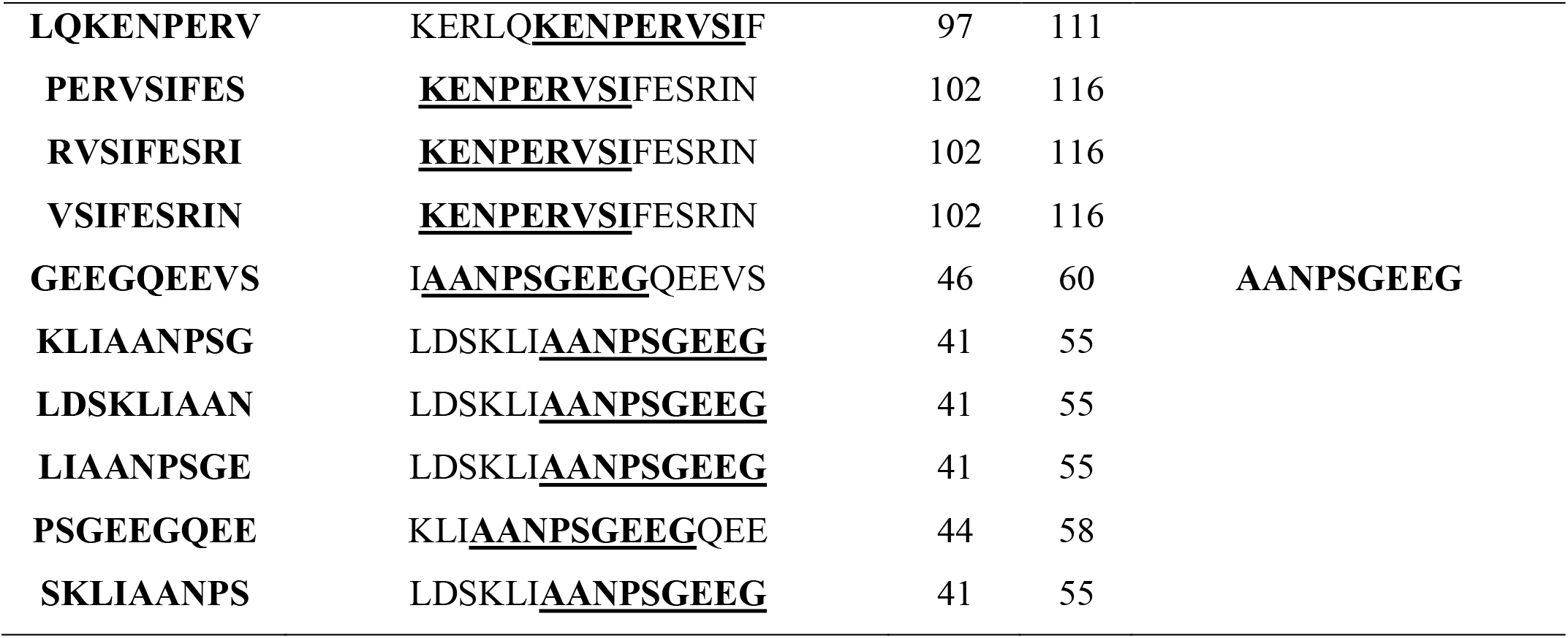
Overlapping between MHC class I and II epitopes.

**Figure 9.**
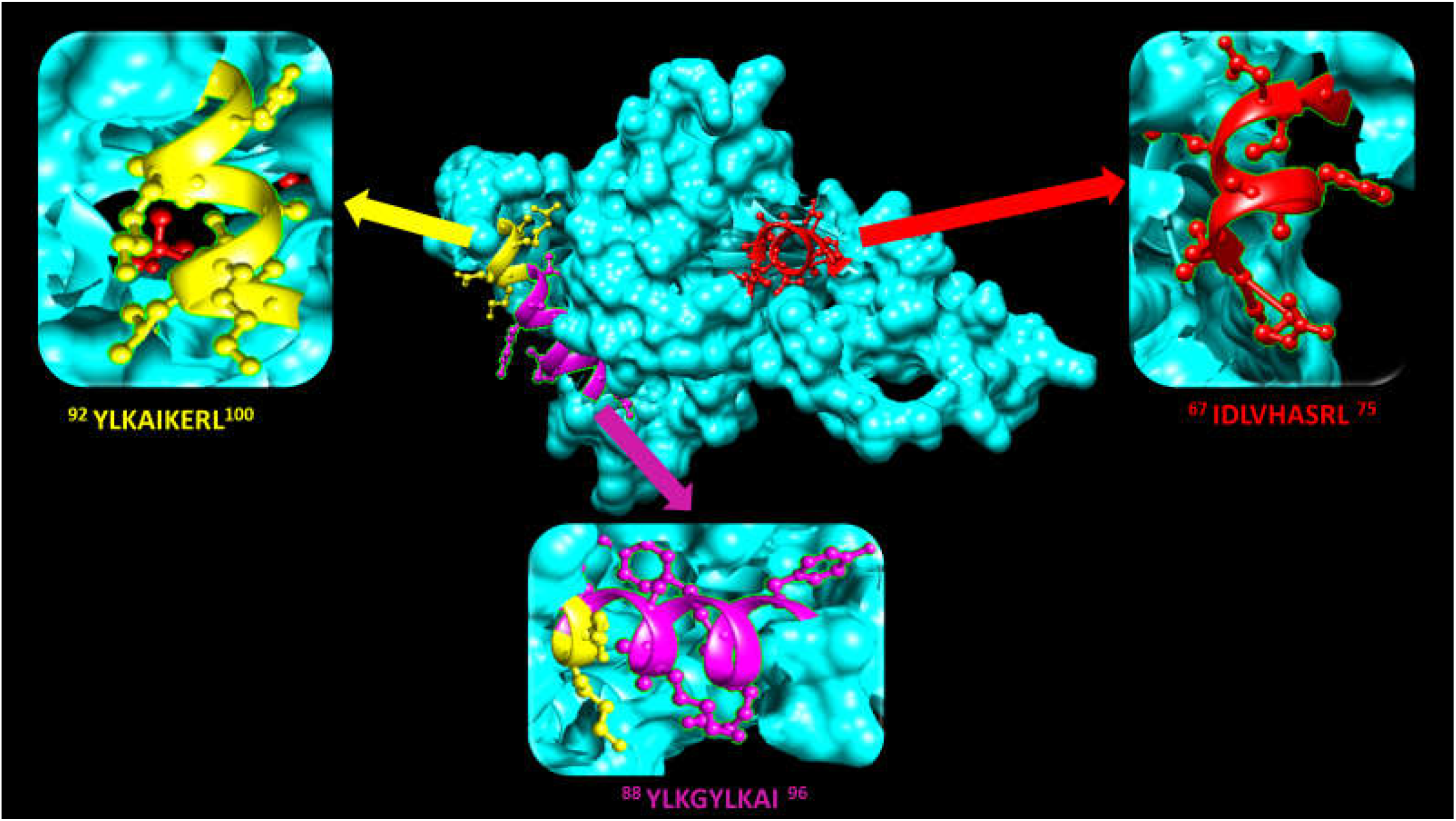
Position of proposed T cell epitopes of *S. japonicum TCTP* that interact with MHC II in structural level using UCSF Chimera (version 1.10.2).

### Population coverage analysis

All predicted T cell epitopes that are interacted with MHC I and MHC II were used for population coverage analysis against the whole world population. The results of population coverage of all epitopes are listed in **Table 6** and **7**. Proposed epitopes which interacted with large number of HLA class I and class II alleles were analyzed against the whole world population, northeast Asia, and southeast Asia. Results of that analysis are presented in **Table 8** and **9**.

**Table 6.**
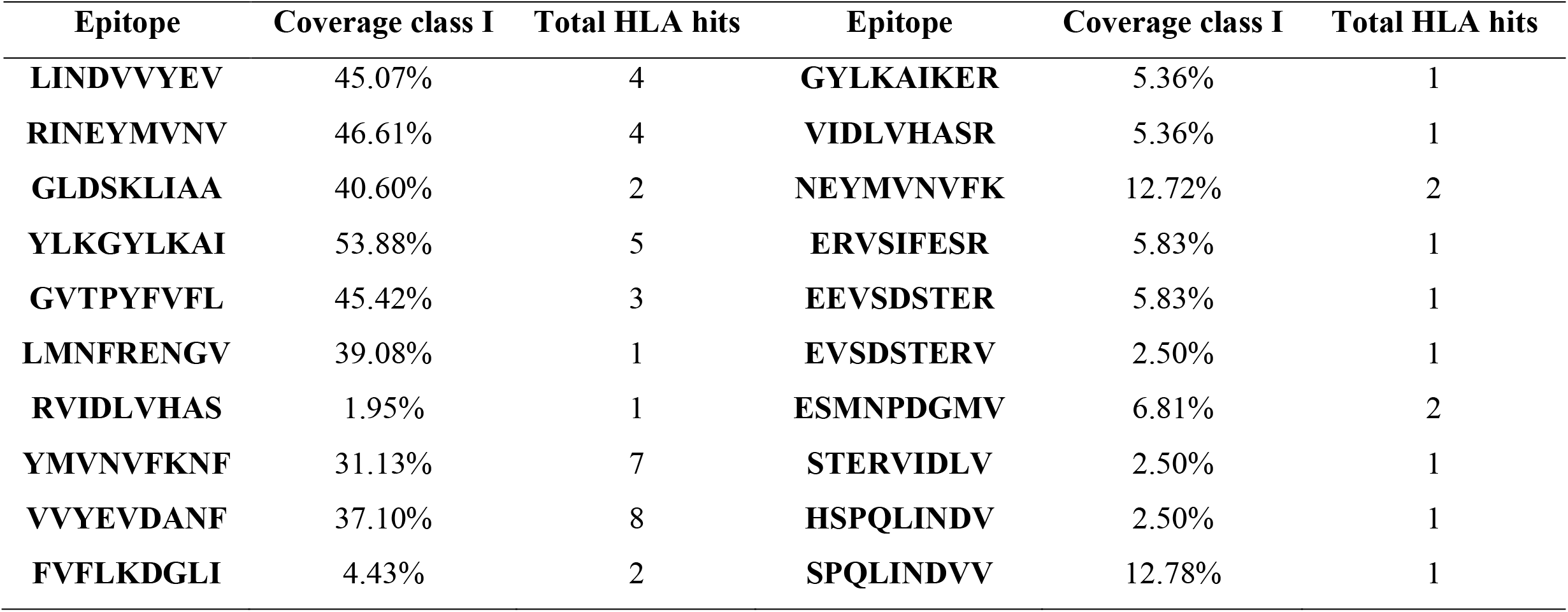
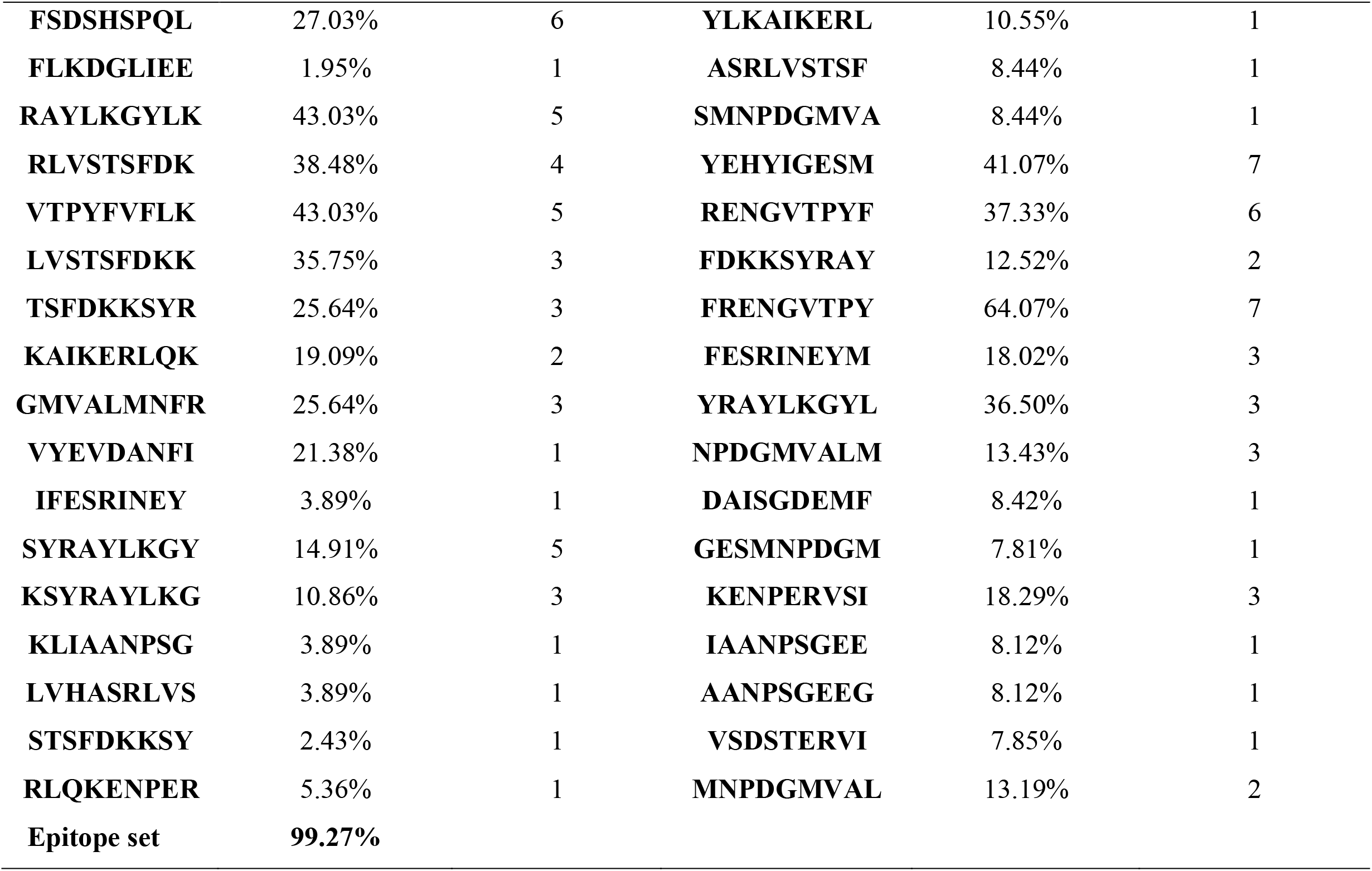
Population coverage of all epitopes in MHC class I in the world.

**Table 7.**
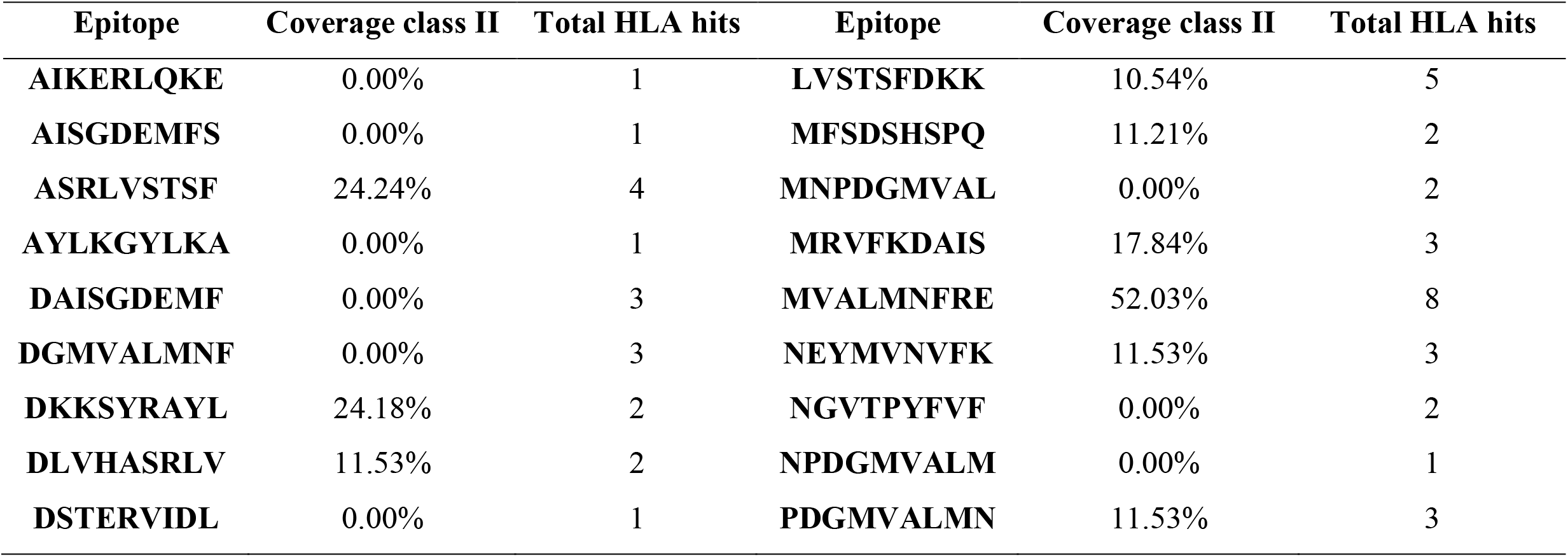
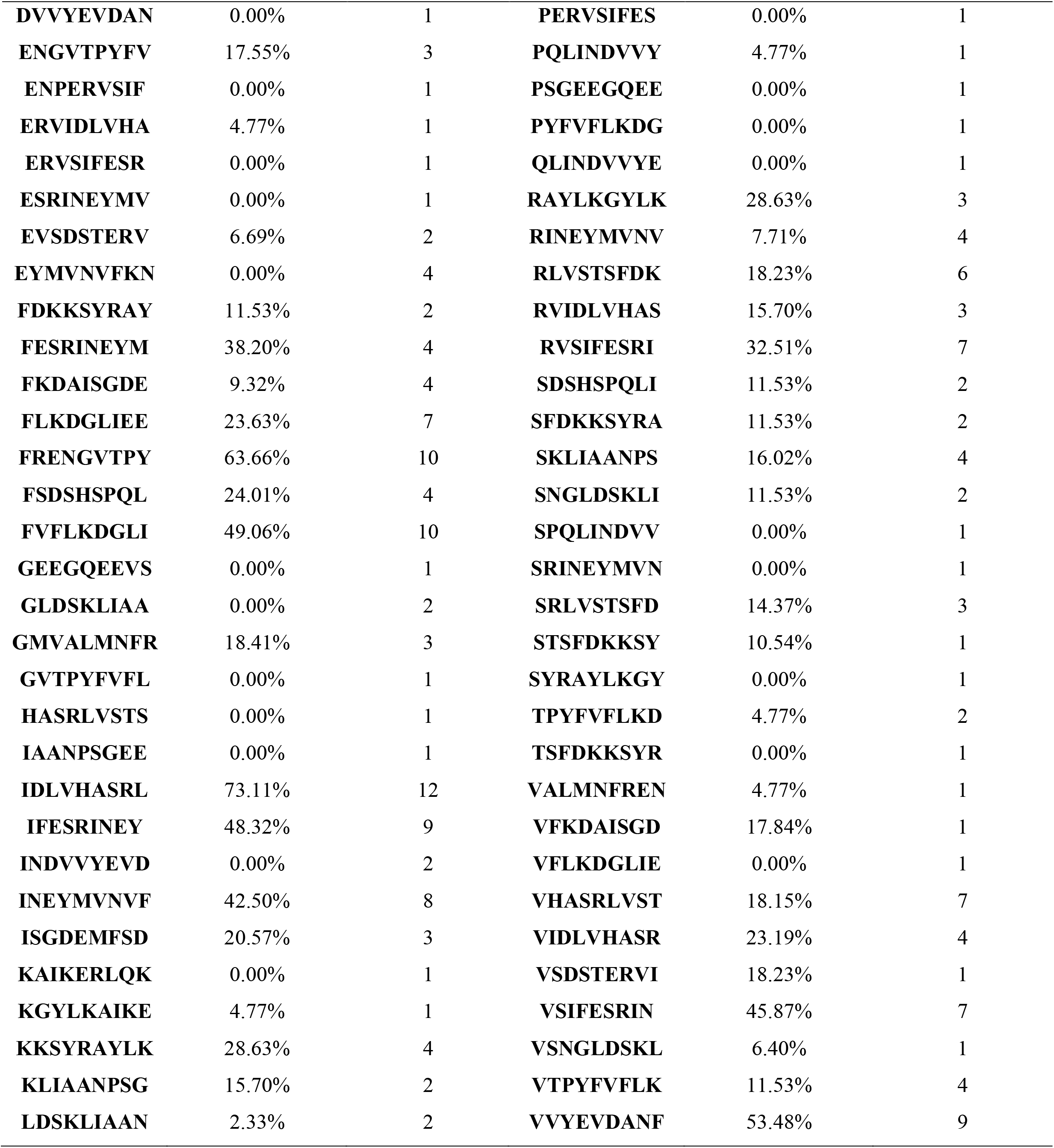
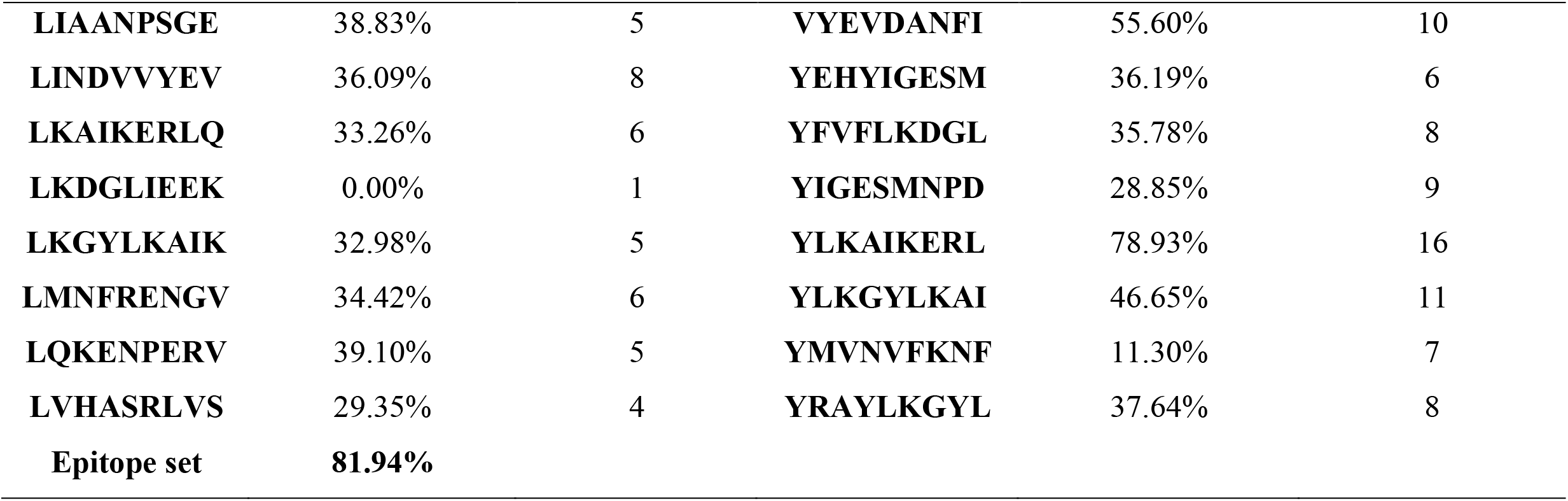
Population coverage of all epitopes in MHC class II in the world.

**Table 8.**
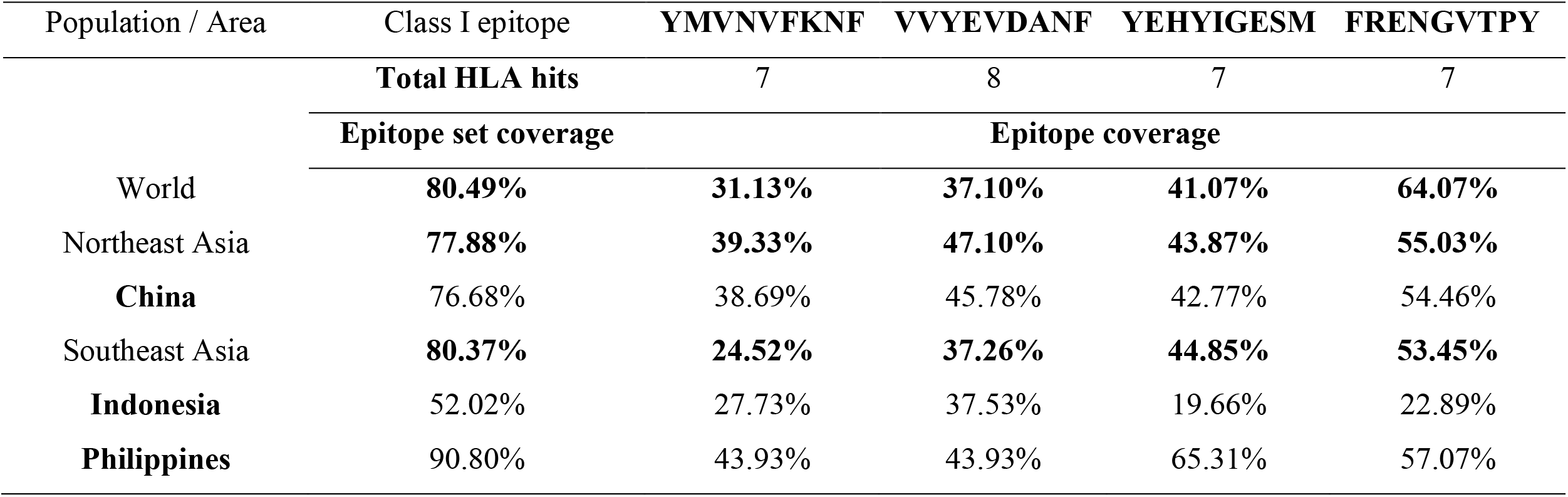
Population coverage of proposed MHC I epitopes in the world, northeast Asia and Southeast Asia.

**Table 9.**
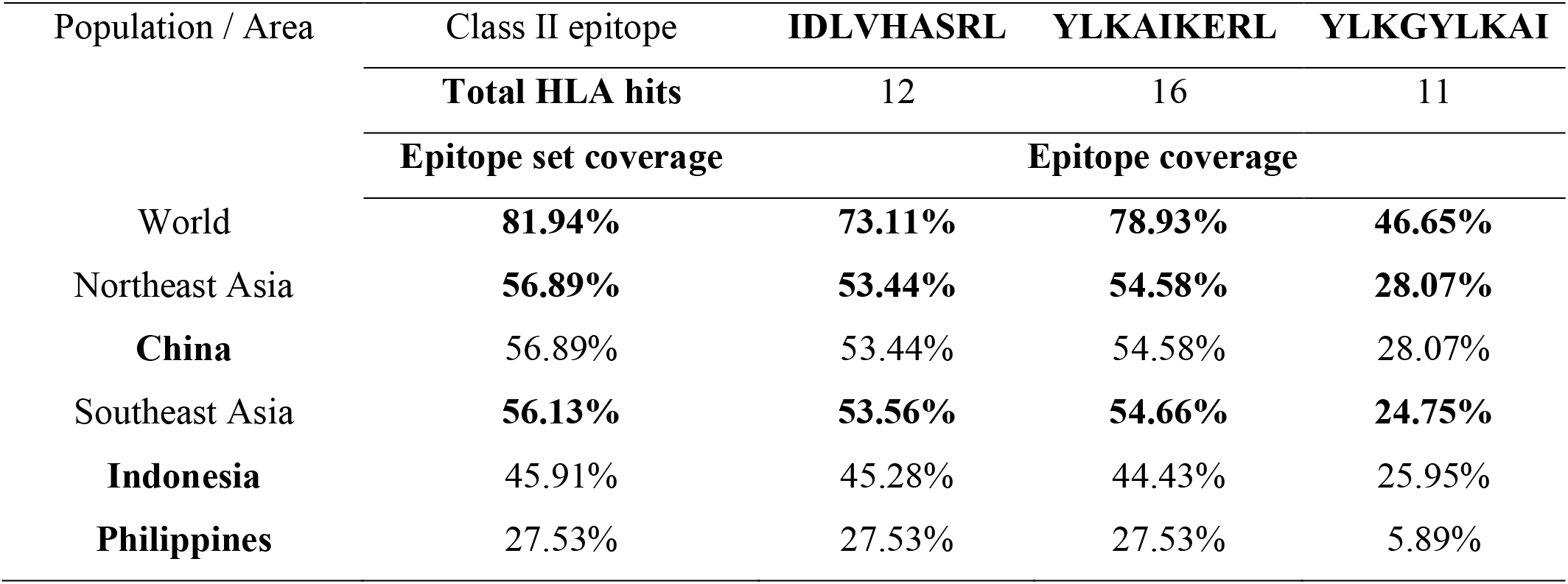
Population coverage of proposed MHC II epitopes in the world, northeast Asia and Southeast Asia.

### Allergenicity test

Allergenicity of both B cell and T cell selected epitopes were tested using AllerTopv.2.0 software to avoid production of IgE antibodies as possible. Results of this test are listed in **Table 10.**

**Table 10.**
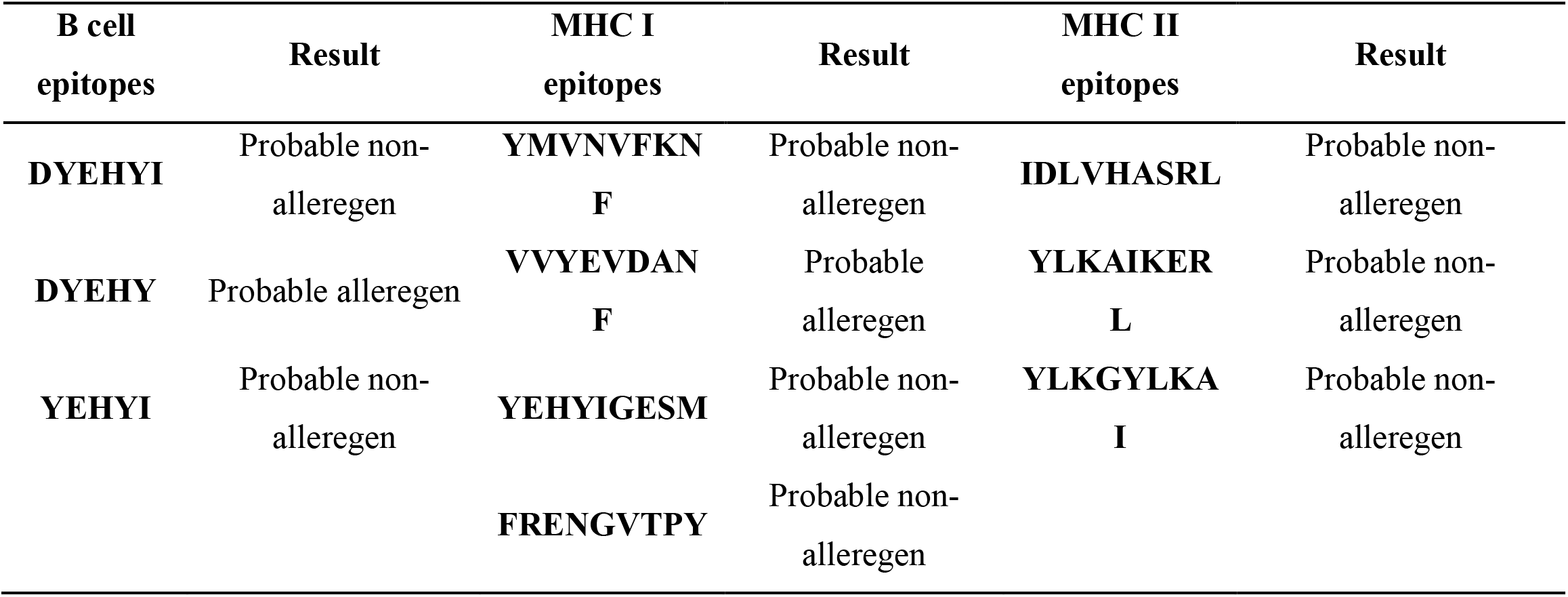
Result of Allergenicity Test of predicted B cell and MHC class I & II epitopes.

## Discussion

There is an urgent need for developing a vaccine as a major aspect to control schistosomiasis as is still making significant public health concern in their endemic countries even with efforts done to control it (20, 63, 64). Therefore, in the current study, we have selected *TCTP* as a target for vaccine design using in silico study, as it has many advantages over traditional peptide vaccine development methods (65–67). Additionally, this in silico approach was used to design peptide vaccine against several diseases and numerous promising peptides vaccine were investigated (68, 69). Bioinformatics analysis of *TCTP* from *Spirometra mansoni* which has the closest evolutionary position with *Schistosoma japonicum*, and *Schistosoma mansoni* revealed that it may be used as a potential vaccine candidate (70). Moreover, promising peptides were predicted from *Madurella mycetomatis TCTP* using an immmunoinformatics approach (71). Besides, the potential role of *TCTP* as a vaccine was investigated by MacDonald SM et al. in 2001 using *Plasmodium falciparum TCTP*. Accordingly, this protein can stimulate histamine release from basophils and IL-8 secretion from eosinophils in vitro and may affect host immune responses in vivo. Additionally, according to Taylor et al. in 2015 *TCTP* from different Plasmodium species resulted in a significant reduction in parasitemia in the early stage of infection in BALB/c mice (72, 73). A set of 22 *TCTP* sequences ‘most of them were collected from China’ were retrieved as FASTA format from NCBI for this study until 9^th^ of December 2017 and conserved regions among them were identified. These strains had high degree of conservancy among them and promiscuous epitopes were selected to be effective antigen for both B cells and T cells.

As B cells play a critical role in adaptive immunity, *SjTCTP* was subjected to Bepipred linear epitope prediction, Emini surface accessibility, and Kolaskar and Tongaonkar antigenicity prediction methods in IEDB to determine their potential effect as B cell antigen. Accordingly, several conserved epitopes having the length ranged from 2 to 19 amino acids were predicted, but not all of them got scores above the threshold in both Emini surface accessibility, and Kolaskar and Tongaonkar antigenicity. However, when we decreased the length of the linear epitope ^**128**^**DYEHYIGESMNPDG**^**141**^ from 14 amino acid to 6, and 5 amino acid, the antigenicity, and surface accessibility score were raised above the threshold values as follow: epitope **YEHYI** from 129 to 133 was found to have one of the highest score in antigenicity (**1.086**) and good score in surface accessibility (**1.054**), followed by **DYEHYI** from 128 to 133 and then **DYEHY** from 128 to 132 which produced the highest score in surface accessibility (**2.511**). On the other hand, T cells recognize epitopes when they are presented by MHC class I and MHC class II. Therefore, binding of peptide and MHC is an important step in the prediction of T-cell epitopes (74). A total of 54 conserved peptides of TCTP were predicted to bind with MHC class I alleles. Four epitopes, **FRENGVTPY**, **YEHYIGESM**, **VVYEVDANF**, and **YMVNVFKNF** were found to bind with a large number of MHC I with high and intermediate affinity. Among four epitopes, epitope ^**26**^**VVYEVDANF**^**34**^had higher affinity to interact with eight alleles, while other epitopes had an affinity to interact with seven alleles of each. In the case of MHC II, a set of 96 conserved epitopes were predicted. Among them, the following epitopes: **YLKAIKERL**, **IDLVHASRL**, and **YLKGYLKAI** were detected to bind with a large number of MHC II alleles. Epitope ^**92**^**YLKAIKERL**^**100**^ was found to interact with the highest number of alleles, sixteen alleles. Epitopes **IDLVHASRL** and **YLKGYLKAI** interacted with twelve and eleven alleles respectively. In addition to that, several overlaps between MHC class I and II were observed in T cell epitopes. These overlaps might increase the possibility of epitopes presentation to T cells via binding to both MHC class I and II.

MHC are highly polymorphic genes, and different populations express different types of MHC alleles. Thus, calculation of population coverage is highly recommended, and the high value of population coverage is required in the development of a universal epitope-based vaccine (57, 74). In this study, the population coverage of all MHC I epitopes in the world is 99.27%, while for MHC II is 81.94% as some alleles were missed from IEDB database and hence did not include in the calculation of population coverage of all predicted MHC II epitopes. Regarding our proposed MHC I and MHC II epitopes, maximum coverage in the world of MHC I (**64.07%**) resulted from epitope **FRENGVTPY** rather than **VVYEVDANF** despite it bound to seven alleles while the later one bound to eight alleles. On the other hand, the maximum population coverage of MHC II (**78.93%**) resulted from **YLKAIKERL** that bound to the largest number of MHC II alleles. When we calculated the population coverage of the proposed MHC I and MHC II epitopes in China, Indonesia, and Philippines ‘the place where *S. japonicum* is endemic’ different coverage of each epitope resulted in each country as there was a difference in the frequency distribution of human leukocyte antigen (HLA) class I and II alleles in those countries (75, 76). In the case of MHC I Epitopes, **FRENGVTPY** is a most promising one in China with population coverage 54.46%, while epitopes **VVYEVDANF** with population coverage 37.53%, and **YEHYIGESM** with coverage 65.31% are the most promising epitopes in Indonesia and Philippines respectively. Whereas, in MHC II epitopes, **YLKAIKERL** with coverage 54.58%, **IDLVHASRL** with coverage 45.28% and both **IDLVHASRL** and **YLKAIKERL** with population coverage 27.53% for each are the most promising epitopes in China, Indonesia, and Philippines respectively.

Although vaccination is an efficient method to prevent the diseases, the potential risk of allergic reactions exists. These allergic reactions are mediated by the reaction of IgE antibody with a vaccine itself or the vaccine components (77–79). Thus, our predicted B and T cell epitopes were subjected to allergenicity test using AllerTopv.2.0. In the case of B cell, two epitopes **DYEHYI** and **YEHYI** have a probability to be a real epitope, while all of the predicted T cell epitopes except **VVYEVDANF** could also be real epitopes as all of them have a non-allergic effect. *Schistosoma japonicum* is still causing a serious problem in its endemic countries, and until now no vaccines have shown a good immunogenic response. Therefore, in this study, our immunoinformatics analysis aided to design a potential immunogenic and safe peptide vaccine which may have a promising preventive ability to control *S. japonicum*. However, to prove the efficacy of the predicted epitopes, additional in vitro and in vivo studies are required along with this computational study.

## Conclusion

Immunoinformatics approaches have enabled the ability to design vaccines avoiding the disadvantages of the conventional methods as they reduce the time and cost required and enhance the safety and efficacy of the predicted epitopes (33, 80–86). Several epitopes were predicted in this study to design a vaccine against *Schistosoma japonicum* using in silico prediction tools. Epitope **YEHYIGESM** is recommended for further in vitro and in vivo studies to confirm its efficacy as a peptide vaccine, as this epitope is shared between two classes of MHC alleles and overlapped with the most promising B cell epitope.

## Acknowledgement

The authors are thankful to all members of Africa City of Technology for their help.

