## Supplemental table 1 for "Translationally controlled tumor protein *TCTP* as Peptide Vaccine against *Schistosoma japonicum*: an immunoinformatics approach"

B cell epitopes:

| Start | End | Peptide | length | Emini surface Score | Antigenicity score |
|---|---|---|---|---|---|
| 7 | 22 | **AISGDEMFSDSHSPQL** | 16 | 0.409 | 1.005 |
| 7 | 21 | **AISGDEMFSDSHSPQ** | 15 | 0.651 | 0.988 |
| 8 | 22 | **ISGDEMFSDSHSPQL** | 15 | 0.531 | 1.001 |
| 7 | 20 | **AISGDEMFSDSHSP** | 14 | 0.477 | 0.986 |
| 8 | 21 | **ISGDEMFSDSHSPQ** | 14 | 0.818 | 0.983 |
| 9 | 22 | **SGDEMFSDSHSPQL** | 14 | 0.962 | 0.99 |
| 7 | 19 | **AISGDEMFSDSHS** | 13 | 0.397 | 0.98 |
| 8 | 20 | **ISGDEMFSDSHSP** | 13 | 0.608 | 0.98 |
| 9 | 21 | **SGDEMFSDSHSPQ** | 13 | 1.502 | 0.97 |
| 10 | 22 | **GDEMFSDSHSPQL** | 13 | 0.924 | 0.988 |
| 7 | 18 | **AISGDEMFSDSH** | 12 | 0.383 | 0.978 |
| 8 | 19 | **ISGDEMFSDSHS** | 12 | 0.508 | 0.973 |
| 9 | 20 | **SGDEMFSDSHSP** | 12 | 1.121 | 0.966 |
| 10 | 21 | **GDEMFSDSHSPQ** | 12 | 1.449 | 0.966 |
| 11 | 22 | **DEMFSDSHSPQL** | 12 | 1.207 | 0.998 |
| 7 | 17 | **AISGDEMFSDS** | 11 | 0.359 | 0.966 |
| 8 | 18 | **ISGDEMFSDSH** | 11 | 0.484 | 0.97 |
| 9 | 19 | **SGDEMFSDSHS** | 11 | 0.925 | 0.957 |
| 10 | 20 | **GDEMFSDSHSP** | 11 | 1.067 | 0.962 |
| 11 | 21 | **DEMFSDSHSPQ** | 11 | 1.868 | 0.975 |
| 12 | 22 | **EMFSDSHSPQL** | 11 | 0.922 | 1.009 |
| 7 | 16 | **AISGDEMFSD** | 10 | 0.346 | 0.961 |
| 8 | 17 | **ISGDEMFSDS** | 10 | 0.459 | 0.956 |
| 9 | 18 | **SGDEMFSDSH** | 10 | 0.891 | 0.952 |
| 10 | 19 | **GDEMFSDSHS** | 10 | 0.891 | 0.952 |
| 11 | 20 | **DEMFSDSHSP** | 10 | 1.392 | 0.971 |
| 12 | 21 | **EMFSDSHSPQ** | 10 | 1.444 | 0.985 |
| 13 | 22 | **MFSDSHSPQL** | 10 | 0.687 | 1.025 |
| 7 | 15 | **AISGDEMFS** | 9 | 0.271 | 0.972 |
| 8 | 16 | **ISGDEMFSD** | 9 | 0.448 | 0.95 |
| 9 | 17 | **SGDEMFSDS** | 9 | 0.856 | 0.934 |
| 10 | 18 | **GDEMFSDSH** | 9 | 0.869 | 0.945 |
| 11 | 19 | **DEMFSDSHS** | 9 | 1.177 | 0.96 |
| 12 | 20 | **EMFSDSHSP** | 9 | 1.09 | 0.982 |
| 13 | 21 | **MFSDSHSPQ** | 9 | 1.09 | 1 |
| 14 | 22 | **FSDSHSPQL** | 9 | 0.908 | 1.047 |
| 7 | 14 | **AISGDEMF** | 8 | 0.267 | 0.967 |
| 8 | 15 | **ISGDEMFS** | 8 | 0.354 | 0.96 |
| 9 | 16 | **SGDEMFSD** | 8 | 0.843 | 0.925 |
| 10 | 17 | **GDEMFSDS** | 8 | 0.843 | 0.925 |

| | | | | | |
|---|---|---|---|---|---|
| 11 | 18 | **DEMFSDSH** | 8 | 1.159 | 0.954 |
| 12 | 19 | **EMFSDSHS** | 8 | 0.93 | 0.972 |
| 13 | 20 | **MFSDSHSP** | 8 | 0.831 | 0.999 |
| 14 | 21 | **FSDSHSPQ** | 8 | 1.454 | 1.022 |
| 15 | 22 | **SDSHSPQL** | 8 | 1.384 | 1.042 |
| 7 | 13 | **AISGDEM** | 7 | 0.403 | 0.949 |
| 8 | 14 | **ISGDEMF** | 7 | 0.346 | 0.953 |
| 9 | 15 | **SGDEMFS** | 7 | 0.661 | 0.933 |
| 10 | 16 | **GDEMFSD** | 7 | 0.823 | 0.912 |
| 11 | 17 | **DEMFSDS** | 7 | 1.115 | 0.932 |
| 12 | 18 | **EMFSDSH** | 7 | 0.908 | 0.966 |
| 13 | 19 | **MFSDSHS** | 7 | 0.703 | 0.989 |
| 14 | 20 | **FSDSHSP** | 7 | 1.098 | 1.023 |
| 15 | 21 | **SDSHSPQ** | 7 | 2.196 | 1.012 |
| 16 | 22 | **DSHSPQL** | 7 | 1.352 | 1.046 |
| 7 | 12 | **AISGDE** | 6 | 0.53 | 0.97 |
| 8 | 13 | **ISGDEM** | 6 | 0.519 | 0.93 |
| 9 | 14 | **SGDEMF** | 6 | 0.641 | 0.92 |
| 10 | 15 | **GDEMFS** | 6 | 0.641 | 0.92 |
| 11 | 16 | **DEMFSD** | 6 | 1.082 | 0.919 |
| 12 | 17 | **EMFSDS** | 6 | 0.868 | 0.943 |
| 13 | 18 | **MFSDSH** | 6 | 0.682 | 0.985 |
| 14 | 19 | **FSDSHS** | 6 | 0.923 | 1.016 |
| 15 | 20 | **SDSHSP** | 6 | 1.649 | 1.012 |
| 16 | 21 | **DSHSPQ** | 6 | 2.131 | 1.012 |
| 17 | 22 | **SHSPQL** | 6 | 1.052 | 1.076 |
| 7 | 11 | **AISGD** | 5 | 0.408 | 0.994 |
| 8 | 12 | **ISGDE** | 5 | 0.699 | 0.951 |
| 9 | 13 | **SGDEM** | 5 | 0.987 | 0.886 |
| 10 | 14 | **GDEMF** | 5 | 0.638 | 0.902 |
| 11 | 15 | **DEMFS** | 5 | 0.863 | 0.929 |
| 12 | 16 | **EMFSD** | 5 | 0.863 | 0.929 |
| 13 | 17 | **MFSDS** | 5 | 0.668 | 0.961 |
| 14 | 18 | **FSDSH** | 5 | 0.919 | 1.017 |
| 15 | 19 | **SDSHS** | 5 | 1.422 | 1.001 |
| 16 | 20 | **DSHSP** | 5 | 1.64 | 1.012 |
| 17 | 21 | **SHSPQ** | 5 | 1.701 | 1.042 |
| 18 | 22 | **HSPQL** | 5 | 1.047 | 1.089 |
| 7 | 10 | **AISG** | 4 | 0.32 | 1.026 |
| 8 | 11 | **ISGD** | 4 | 0.529 | 0.976 |
| 9 | 12 | **SGDE** | 4 | 1.306 | 0.901 |
| 10 | 13 | **GDEM** | 4 | 0.964 | 0.854 |
| 11 | 14 | **DEMF** | 4 | 0.844 | 0.909 |

| | | | | | |
|---|---|---|---|---|---|
| 12 | 15 | **EMFS** | 4 | 0.677 | 0.945 |
| 13 | 16 | **MFSD** | 4 | 0.653 | 0.949 |
| 14 | 17 | **FSDS** | 4 | 0.884 | 0.995 |
| 15 | 18 | **SDSH** | 4 | 1.389 | 0.999 |
| 16 | 19 | **DSHS** | 4 | 1.389 | 0.999 |
| 17 | 20 | **SHSP** | 4 | 1.286 | 1.048 |
| 18 | 21 | **HSPQ** | 4 | 1.662 | 1.049 |
| 19 | 22 | **SPQL** | 4 | 1.008 | 1.085 |
| 7 | 9 | **AIS** | 3 | 0.415 | 1.076 |
| 8 | 10 | **ISG** | 3 | 0.407 | 1.013 |
| 9 | 11 | **SGD** | 3 | 0.969 | 0.917 |
| 10 | 12 | **GDE** | 3 | 1.252 | 0.864 |
| 11 | 13 | **DEM** | 3 | 1.252 | 0.848 |
| 12 | 14 | **EMF** | 3 | 0.649 | 0.923 |
| 13 | 15 | **MFS** | 3 | 0.503 | 0.976 |
| 14 | 16 | **FSD** | 3 | 0.848 | 0.99 |
| 15 | 17 | **SDS** | 3 | 1.312 | 0.963 |
| 16 | 18 | **DSH** | 3 | 1.333 | 0.994 |
| 17 | 19 | **SHS** | 3 | 1.069 | 1.043 |
| 18 | 20 | **HSP** | 3 | 1.234 | 1.06 |
| 19 | 21 | **SPQ** | 3 | 1.57 | 1.03 |
| 20 | 22 | **PQL** | 3 | 0.966 | 1.11 |
| 39 | 41 | **NGL** | 3 | 0.574 | 0.967 |
| 46 | 64 | **IAANPSGEEGQEEVSDSTE** | 19 | 1.4 | 0.965 |
| 46 | 63 | **IAANPSGEEGQEEVSDST** | 18 | 1.014 | 0.971 |
| 47 | 64 | **AANPSGEEGQEEVSDSTE** | 18 | 2.505 | 0.954 |
| 46 | 62 | **IAANPSGEEGQEEVSDS** | 17 | 0.909 | 0.975 |
| 47 | 63 | **AANPSGEEGQEEVSDST** | 17 | 1.871 | 0.961 |
| 48 | 64 | **ANPSGEEGQEEVSDSTE** | 17 | 3.208 | 0.948 |
| 46 | 61 | **IAANPSGEEGQEEVSD** | 16 | 0.878 | 0.972 |
| 47 | 62 | **AANPSGEEGQEEVSDS** | 16 | 1.679 | 0.964 |
| 48 | 63 | **ANPSGEEGQEEVSDST** | 16 | 2.399 | 0.954 |
| 49 | 64 | **NPSGEEGQEEVSDSTE** | 16 | 4.112 | 0.941 |
| 46 | 60 | **IAANPSGEEGQEEVS** | 15 | 0.691 | 0.98 |
| 47 | 61 | **AANPSGEEGQEEVSD** | 15 | 1.646 | 0.961 |
| 48 | 62 | **ANPSGEEGQEEVSDS** | 15 | 2.183 | 0.957 |
| 49 | 63 | **NPSGEEGQEEVSDST** | 15 | 3.119 | 0.947 |
| 50 | 64 | **PSGEEGQEEVSDSTE** | 15 | 3.359 | 0.952 |
| 46 | 59 | **IAANPSGEEGQEEV** | 14 | 0.654 | 0.977 |
| 47 | 60 | **AANPSGEEGQEEVS** | 14 | 1.251 | 0.967 |
| 48 | 61 | **ANPSGEEGQEEVSD** | 14 | 2.068 | 0.953 |
| 49 | 62 | **NPSGEEGQEEVSDS** | 14 | 2.743 | 0.949 |
| 50 | 63 | **PSGEEGQEEVSDST** | 14 | 2.461 | 0.959 |

| | | | | | |
|---|---|---|---|---|---|
| 51 | 64 | **SGEEGQEEVSDSTE** | 14 | 2.757 | 0.944 |
| 46 | 58 | **IAANPSGEEGQEE** | 13 | 1.135 | 0.946 |
| 47 | 59 | **AANPSGEEGQEEV** | 13 | 1.201 | 0.964 |
| 48 | 60 | **ANPSGEEGQEEVS** | 13 | 1.594 | 0.96 |
| 49 | 61 | **NPSGEEGQEEVSD** | 13 | 2.634 | 0.945 |
| 50 | 62 | **PSGEEGQEEVSDS** | 13 | 2.195 | 0.963 |
| 51 | 63 | **SGEEGQEEVSDST** | 13 | 2.049 | 0.951 |
| 52 | 64 | **GEEGQEEVSDSTE** | 13 | 2.648 | 0.938 |
| 46 | 57 | **IAANPSGEEGQE** | 12 | 0.847 | 0.954 |
| 47 | 58 | **AANPSGEEGQEE** | 12 | 2.093 | 0.929 |
| 48 | 59 | **ANPSGEEGQEEV** | 12 | 1.538 | 0.955 |
| 49 | 60 | **NPSGEEGQEEVS** | 12 | 2.04 | 0.951 |
| 50 | 61 | **PSGEEGQEEVSD** | 12 | 2.118 | 0.959 |
| 51 | 62 | **SGEEGQEEVSDS** | 12 | 1.836 | 0.954 |
| 52 | 63 | **GEEGQEEVSDST** | 12 | 1.977 | 0.946 |
| 53 | 64 | **EEGQEEVSDSTE** | 12 | 3.459 | 0.944 |
| 46 | 56 | **IAANPSGEEGQ** | 11 | 0.624 | 0.963 |
| 47 | 57 | **AANPSGEEGQE** | 11 | 1.542 | 0.936 |
| 48 | 58 | **ANPSGEEGQEE** | 11 | 2.643 | 0.917 |
| 49 | 59 | **NPSGEEGQEEV** | 11 | 1.942 | 0.946 |
| 50 | 60 | **PSGEEGQEEVS** | 11 | 1.618 | 0.967 |
| 51 | 61 | **SGEEGQEEVSD** | 11 | 1.748 | 0.949 |
| 52 | 62 | **GEEGQEEVSDS** | 11 | 1.748 | 0.949 |
| 53 | 63 | **EEGQEEVSDST** | 11 | 2.549 | 0.952 |
| 54 | 64 | **EGQEEVSDSTE** | 11 | 2.549 | 0.952 |
| 46 | 55 | **IAANPSGEEG** | 10 | 0.465 | 0.958 |
| 47 | 56 | **AANPSGEEGQ** | 10 | 1.149 | 0.945 |
| 48 | 57 | **ANPSGEEGQE** | 10 | 1.97 | 0.923 |
| 49 | 58 | **NPSGEEGQEE** | 10 | 3.377 | 0.902 |
| 50 | 59 | **PSGEEGQEEV** | 10 | 1.559 | 0.963 |
| 51 | 60 | **SGEEGQEEVS** | 10 | 1.351 | 0.957 |
| 52 | 61 | **GEEGQEEVSD** | 10 | 1.683 | 0.943 |
| 53 | 62 | **EEGQEEVSDS** | 10 | 2.279 | 0.957 |
| 54 | 63 | **EGQEEVSDST** | 10 | 1.9 | 0.962 |
| 55 | 64 | **GQEEVSDSTE** | 10 | 1.9 | 0.962 |
| 46 | 54 | **IAANPSGEE** | 9 | 0.615 | 0.968 |
| 47 | 55 | **AANPSGEEG** | 9 | 0.868 | 0.937 |
| 48 | 56 | **ANPSGEEGQ** | 9 | 1.487 | 0.931 |
| 49 | 57 | **NPSGEEGQE** | 9 | 2.55 | 0.908 |
| 50 | 58 | **PSGEEGQEE** | 9 | 2.746 | 0.916 |
| 51 | 59 | **SGEEGQEEV** | 9 | 1.318 | 0.951 |
| 52 | 60 | **GEEGQEEVS** | 9 | 1.318 | 0.951 |
| 53 | 61 | **EEGQEEVSD** | 9 | 2.224 | 0.95 |

| | | | | | |
|---|---|---|---|---|---|
| 54 | 62 | **EGQEEVSDS** | 9 | 1.721 | 0.968 |
| 55 | 63 | **GQEEVSDST** | 9 | 1.434 | 0.975 |
| 56 | 64 | **QEEVSDSTE** | 9 | 2.51 | 0.972 |
| 46 | 53 | **IAANPSGE** | 8 | 0.468 | 0.982 |
| 47 | 54 | **AANPSGEE** | 8 | 1.157 | 0.945 |
| 48 | 55 | **ANPSGEEG** | 8 | 1.134 | 0.921 |
| 49 | 56 | **NPSGEEGQ** | 8 | 1.943 | 0.915 |
| 50 | 57 | **PSGEEGQE** | 8 | 2.093 | 0.924 |
| 51 | 58 | **SGEEGQEE** | 8 | 2.344 | 0.897 |
| 52 | 59 | **GEEGQEEV** | 8 | 1.298 | 0.944 |
| 53 | 60 | **EEGQEEVS** | 8 | 1.758 | 0.961 |
| 54 | 61 | **EGQEEVSD** | 8 | 1.695 | 0.963 |
| 55 | 62 | **GQEEVSDS** | 8 | 1.312 | 0.983 |
| 56 | 63 | **QEEVSDST** | 8 | 1.913 | 0.987 |
| 57 | 64 | **EEVSDSTE** | 8 | 1.913 | 0.967 |
| 46 | 52 | **IAANPSG** | 7 | 0.354 | 1.001 |
| 47 | 53 | **AANPSGE** | 7 | 0.874 | 0.958 |
| 48 | 54 | **ANPSGEE** | 7 | 1.498 | 0.927 |
| 49 | 55 | **NPSGEEG** | 7 | 1.468 | 0.9 |
| 50 | 56 | **PSGEEGQ** | 7 | 1.581 | 0.934 |
| 51 | 57 | **SGEEGQE** | 7 | 1.77 | 0.904 |
| 52 | 58 | **GEEGQEE** | 7 | 2.288 | 0.881 |
| 53 | 59 | **EEGQEEV** | 7 | 1.716 | 0.954 |
| 54 | 60 | **EGQEEVS** | 7 | 1.328 | 0.977 |
| 55 | 61 | **GQEEVSD** | 7 | 1.28 | 0.979 |
| 56 | 62 | **QEEVSDS** | 7 | 1.734 | 0.999 |
| 57 | 63 | **EEVSDST** | 7 | 1.445 | 0.983 |
| 58 | 64 | **EVSDSTE** | 7 | 1.445 | 0.983 |
| 46 | 51 | **IAANPS** | 6 | 0.465 | 1.022 |
| 47 | 52 | **AANPSG** | 6 | 0.656 | 0.976 |
| 48 | 53 | **ANPSGE** | 6 | 1.125 | 0.94 |
| 49 | 54 | **NPSGEE** | 6 | 1.929 | 0.905 |
| 50 | 55 | **PSGEEG** | 6 | 1.187 | 0.921 |
| 51 | 56 | **SGEEGQ** | 6 | 1.329 | 0.913 |
| 52 | 57 | **GEEGQE** | 6 | 1.718 | 0.886 |
| 53 | 58 | **EEGQEE** | 6 | 3.006 | 0.882 |
| 54 | 59 | **EGQEEV** | 6 | 1.288 | 0.971 |
| 55 | 60 | **GQEEVS** | 6 | 0.997 | 0.998 |
| 56 | 61 | **QEEVSD** | 6 | 1.682 | 0.996 |
| 57 | 62 | **EEVSDS** | 6 | 1.302 | 0.996 |
| 58 | 63 | **EVSDST** | 6 | 1.085 | 1.006 |
| 59 | 64 | **VSDSTE** | 6 | 1.085 | 1.005 |
| 46 | 50 | **IAANP** | 5 | 0.462 | 1.024 |

| | | | | | |
|---|---|---|---|---|---|
| 47 | 51 | **AANPS** | 5 | 0.884 | 0.996 |
| 48 | 52 | **ANPSG** | 5 | 0.866 | 0.958 |
| 49 | 53 | **NPSGE** | 5 | 1.485 | 0.915 |
| 50 | 54 | **PSGEE** | 5 | 1.599 | 0.93 |
| 51 | 55 | **SGEEG** | 5 | 1.023 | 0.892 |
| 52 | 56 | **GEEGQ** | 5 | 1.322 | 0.893 |
| 53 | 57 | **EEGQE** | 5 | 2.314 | 0.888 |
| 54 | 58 | **EGQEE** | 5 | 2.314 | 0.888 |
| 55 | 59 | **GQEEV** | 5 | 0.992 | 0.995 |
| 56 | 60 | **QEEVS** | 5 | 1.343 | 1.022 |
| 57 | 61 | **EEVSD** | 5 | 1.295 | 0.993 |
| 58 | 62 | **EVSDS** | 5 | 1.002 | 1.025 |
| 59 | 63 | **VSDST** | 5 | 0.835 | 1.036 |
| 60 | 64 | **SDSTE** | 5 | 1.948 | 0.93 |
| 46 | 49 | **IAAN** | 4 | 0.392 | 1.014 |
| 47 | 50 | **AANP** | 4 | 0.864 | 0.992 |
| 48 | 51 | **ANPS** | 4 | 1.146 | 0.979 |
| 49 | 52 | **NPSG** | 4 | 1.123 | 0.932 |
| 50 | 53 | **PSGE** | 4 | 1.209 | 0.95 |
| 51 | 54 | **SGEE** | 4 | 1.354 | 0.897 |
| 52 | 55 | **GEEG** | 4 | 1 | 0.863 |
| 53 | 56 | **EEGQ** | 4 | 1.75 | 0.898 |
| 54 | 57 | **EGQE** | 4 | 1.75 | 0.898 |
| 55 | 58 | **GQEE** | 4 | 1.75 | 0.898 |
| 56 | 59 | **QEEV** | 4 | 1.312 | 1.025 |
| 57 | 60 | **EEVS** | 4 | 1.016 | 1.024 |
| 58 | 61 | **EVSD** | 4 | 0.979 | 1.028 |
| 59 | 62 | **VSDS** | 4 | 0.758 | 1.068 |
| 60 | 63 | **SDST** | 4 | 1.474 | 0.95 |
| 61 | 64 | **DSTE** | 4 | 1.904 | 0.909 |
| 46 | 48 | **IAA** | 3 | 0.313 | 1.093 |
| 47 | 49 | **AAN** | 3 | 0.718 | 0.968 |
| 48 | 50 | **ANP** | 3 | 1.099 | 0.968 |
| 49 | 51 | **NPS** | 3 | 1.458 | 0.951 |
| 50 | 52 | **PSG** | 3 | 0.897 | 0.983 |
| 51 | 53 | **SGE** | 3 | 1.005 | 0.912 |
| 52 | 54 | **GEE** | 3 | 1.299 | 0.859 |
| 53 | 55 | **EEG** | 3 | 1.299 | 0.859 |
| 54 | 56 | **EGQ** | 3 | 1.299 | 0.913 |
| 55 | 57 | **GQE** | 3 | 1.299 | 0.913 |
| 56 | 58 | **QEE** | 3 | 2.273 | 0.906 |
| 57 | 59 | **EEV** | 3 | 0.974 | 1.028 |
| 58 | 60 | **EVS** | 3 | 0.754 | 1.082 |

| | | | | | |
|---|---|---|---|---|---|
| 59 | 61 | **VSD** | 3 | 0.727 | 1.087 |
| 60 | 62 | **SDS** | 3 | 1.312 | 0.963 |
| 61 | 63 | **DST** | 3 | 1.413 | 0.929 |
| 62 | 64 | **STE** | 3 | 1.466 | 0.924 |
| 79 | 85 | **SFDKKSY** | 7 | 2.441 | 1 |
| 79 | 84 | **SFDKKS** | 6 | 2.025 | 0.973 |
| 80 | 85 | **FDKKSY** | 6 | 2.368 | 0.998 |
| 79 | 83 | **SFDKK** | 5 | 2.015 | 0.966 |
| 80 | 84 | **FDKKS** | 5 | 2.015 | 0.966 |
| 81 | 85 | **DKKSY** | 5 | 3.645 | 0.98 |
| 79 | 82 | **SFDK** | 4 | 1.319 | 0.975 |
| 80 | 83 | **FDKK** | 4 | 1.969 | 0.954 |
| 81 | 84 | **DKKS** | 4 | 3.047 | 0.934 |
| 82 | 85 | **KKSY** | 4 | 2.859 | 1.008 |
| 79 | 81 | **SFD** | 3 | 0.848 | 0.99 |
| 80 | 82 | **FDK** | 3 | 1.265 | 0.962 |
| 81 | 83 | **DKK** | 3 | 2.923 | 0.909 |
| 82 | 84 | **KKS** | 3 | 2.345 | 0.957 |
| 83 | 85 | **KSY** | 3 | 1.838 | 1.034 |
| 99 | 108 | **RLQKENPERV** | 10 | 4.029 | 0.987 |
| 99 | 107 | **RLQKENPER** | 9 | 7.098 | 0.943 |
| 100 | 108 | **LQKENPERV** | 9 | 2.69 | 0.999 |
| 99 | 106 | **RLQKENPE** | 8 | 4.783 | 0.951 |
| 100 | 107 | **LQKENPER** | 8 | 4.783 | 0.951 |
| 101 | 108 | **QKENPERV** | 8 | 4.304 | 0.968 |
| 99 | 105 | **RLQKENP** | 7 | 3.613 | 0.966 |
| 100 | 106 | **LQKENPE** | 7 | 3.194 | 0.962 |
| 101 | 107 | **QKENPER** | 7 | 7.587 | 0.909 |
| 102 | 108 | **KENPERV** | 7 | 3.252 | 0.961 |
| 99 | 104 | **RLQKEN** | 6 | 3.038 | 0.949 |
| 100 | 105 | **LQKENP** | 6 | 2.398 | 0.981 |
| 101 | 106 | **QKENPE** | 6 | 5.037 | 0.914 |
| 102 | 107 | **KENPER** | 6 | 5.696 | 0.891 |
| 103 | 108 | **ENPERV** | 6 | 2.114 | 0.966 |
| 99 | 103 | **RLQKE** | 5 | 2.518 | 0.984 |
| 100 | 104 | **LQKEN** | 5 | 2.068 | 0.964 |
| 101 | 105 | **QKENP** | 5 | 3.877 | 0.927 |
| 102 | 106 | **KENPE** | 5 | 3.877 | 0.894 |
| 103 | 107 | **ENPER** | 5 | 3.797 | 0.883 |
| 104 | 108 | **NPERV** | 5 | 1.627 | 0.989 |
| 99 | 102 | **RLQK** | 4 | 1.904 | 1.017 |
| 100 | 103 | **LQKE** | 4 | 1.684 | 1.011 |
| 101 | 104 | **QKEN** | 4 | 3.284 | 0.893 |

| | | | | | |
|---|---|---|---|---|---|
| 102 | 105 | **KENP** | 4 | 2.932 | 0.905 |
| 103 | 106 | **ENPE** | 4 | 2.539 | 0.885 |
| 104 | 107 | **NPER** | 4 | 2.871 | 0.891 |
| 105 | 108 | **PERV** | 4 | 1.325 | 1.043 |
| 99 | 101 | **RLQ** | 3 | 1.224 | 1.046 |
| 100 | 102 | **LQK** | 3 | 1.25 | 1.065 |
| 101 | 103 | **QKE** | 3 | 2.625 | 0.932 |
| 102 | 104 | **KEN** | 3 | 2.437 | 0.852 |
| 103 | 105 | **ENP** | 3 | 1.884 | 0.897 |
| 104 | 106 | **NPE** | 3 | 1.884 | 0.897 |
| 105 | 107 | **PER** | 3 | 2.295 | 0.929 |
| 106 | 108 | **ERV** | 3 | 1.102 | 1.036 |
| 128 | 141 | **DYEHYIGESMNPDG** | 14 | 1.533 | 0.96 |
| 128 | 140 | **DYEHYIGESMNPD** | 13 | 1.994 | 0.967 |
| 129 | 141 | **YEHYIGESMNPDG** | 13 | 1.182 | 0.967 |
| 128 | 139 | **DYEHYIGESMNP** | 12 | 1.544 | 0.975 |
| 129 | 140 | **YEHYIGESMNPD** | 12 | 1.544 | 0.975 |
| 130 | 141 | **EHYIGESMNPDG** | 12 | 0.975 | 0.951 |
| 128 | 138 | **DYEHYIGESMN** | 11 | 1.274 | 0.967 |
| 129 | 139 | **YEHYIGESMNP** | 11 | 1.18 | 0.985 |
| 130 | 140 | **EHYIGESMNPD** | 11 | 1.257 | 0.958 |
| 131 | 141 | **HYIGESMNPDG** | 11 | 0.718 | 0.96 |
| 128 | 137 | **DYEHYIGESM** | 10 | 1.023 | 0.986 |
| 129 | 138 | **YEHYIGESMN** | 10 | 0.985 | 0.977 |
| 130 | 139 | **EHYIGESMNP** | 10 | 0.972 | 0.967 |
| 131 | 140 | **HYIGESMNPD** | 10 | 0.937 | 0.969 |
| 132 | 141 | **YIGESMNPDG** | 10 | 0.681 | 0.946 |
| 128 | 136 | **DYEHYIGES** | 9 | 1.351 | 1.004 |
| 129 | 137 | **YEHYIGESM** | 9 | 0.801 | 0.999 |
| 130 | 138 | **EHYIGESMN** | 9 | 0.822 | 0.956 |
| 131 | 139 | **HYIGESMNP** | 9 | 0.734 | 0.98 |
| 132 | 140 | **YIGESMNPD** | 9 | 0.9 | 0.954 |
| 133 | 141 | **IGESMNPDG** | 9 | 0.569 | 0.922 |
| 128 | 135 | **DYEHYIGE** | 8 | 1.331 | 1.003 |
| 129 | 136 | **YEHYIGES** | 8 | 1.068 | 1.021 |
| 130 | 137 | **EHYIGESM** | 8 | 0.674 | 0.979 |
| 131 | 138 | **HYIGESMN** | 8 | 0.626 | 0.97 |
| 132 | 139 | **YIGESMNP** | 8 | 0.712 | 0.965 |
| 133 | 140 | **IGESMNPD** | 8 | 0.758 | 0.928 |
| 134 | 141 | **GESMNPDG** | 8 | 1.071 | 0.893 |
| 128 | 134 | **DYEHYIG** | 7 | 1.005 | 1.024 |
| 129 | 135 | **YEHYIGE** | 7 | 1.042 | 1.022 |
| 130 | 136 | **EHYIGES** | 7 | 0.892 | 1.001 |

| | | | | | |
|---|---|---|---|---|---|
| 131 | 137 | **HYIGESM** | 7 | 0.509 | 0.997 |
| 132 | 138 | **YIGESMN** | 7 | 0.602 | 0.95 |
| 133 | 139 | **IGESMNP** | 7 | 0.594 | 0.936 |
| 134 | 140 | **GESMNPD** | 7 | 1.415 | 0.896 |
| 135 | 141 | **ESMNPDG** | 7 | 1.415 | 0.896 |
| 128 | 133 | ***DYEHYI** | 6 | 1.321 | 1.049 |
| 129 | 134 | **YEHYIG** | 6 | 0.783 | 1.051 |
| 130 | 135 | **EHYIGE** | 6 | 0.865 | 0.999 |
| 131 | 136 | **HYIGES** | 6 | 0.669 | 1.026 |
| 132 | 137 | **YIGESM** | 6 | 0.487 | 0.979 |
| 133 | 138 | **IGESMN** | 6 | 0.5 | 0.915 |
| 134 | 139 | **GESMNP** | 6 | 1.102 | 0.901 |
| 135 | 140 | **ESMNPD** | 6 | 1.86 | 0.899 |
| 136 | 141 | **SMNPDG** | 6 | 1.063 | 0.903 |
| 128 | 132 | ***DYEHY** | 5 | 2.511 | 1.029 |
| 129 | 133 | ***YEHYI** | 5 | 1.054 | 1.086 |
| 130 | 134 | **EHYIG** | 5 | 0.666 | 1.029 |
| 131 | 135 | **HYIGE** | 5 | 0.666 | 1.029 |
| 132 | 136 | **YIGES** | 5 | 0.656 | 1.01 |
| 133 | 137 | **IGESM** | 5 | 0.414 | 0.943 |
| 134 | 138 | **GESMN** | 5 | 0.95 | 0.868 |
| 135 | 139 | **ESMNP** | 5 | 1.485 | 0.906 |
| 136 | 140 | **SMNPD** | 5 | 1.431 | 0.909 |
| 137 | 141 | **MNPDG** | 5 | 1.057 | 0.881 |
| 128 | 131 | **DYEH** | 4 | 2.099 | 0.996 |
| 129 | 132 | **YEHY** | 4 | 1.97 | 1.07 |
| 130 | 133 | **EHYI** | 4 | 0.881 | 1.067 |
| 131 | 134 | **HYIG** | 4 | 0.504 | 1.073 |
| 132 | 135 | **YIGE** | 4 | 0.641 | 1.01 |
| 133 | 136 | **IGES** | 4 | 0.548 | 0.972 |
| 134 | 137 | **GESM** | 4 | 0.774 | 0.891 |
| 135 | 138 | **ESMN** | 4 | 1.257 | 0.866 |
| 136 | 139 | **SMNP** | 4 | 1.123 | 0.919 |
| 137 | 140 | **MNPD** | 4 | 1.399 | 0.883 |
| 138 | 141 | **NPDG** | 4 | 1.399 | 0.895 |
| 128 | 130 | **DYE** | 3 | 1.983 | 0.959 |
| 129 | 131 | **YEH** | 3 | 1.616 | 1.039 |
| 130 | 132 | **EHY** | 3 | 1.616 | 1.039 |
| 131 | 133 | **HYI** | 3 | 0.654 | 1.139 |
| 132 | 134 | **YIG** | 3 | 0.476 | 1.062 |
| 133 | 135 | **IGE** | 3 | 0.526 | 0.959 |
| 134 | 136 | **GES** | 3 | 1.005 | 0.912 |
| 135 | 137 | **ESM** | 3 | 1.005 | 0.896 |

| | | | | | |
|---|---|---|---|---|---|
| 136 | 138 | **SMN** | 3 | 0.933 | 0.871 |
| 137 | 139 | **MNP** | 3 | 1.077 | 0.889 |
| 138 | 140 | **NPD** | 3 | 1.817 | 0.902 |
| 139 | 141 | **PDG** | 3 | 1.118 | 0.935 |
| 150 | 154 | **ENGVT** | 5 | 0.767 | 0.959 |
| 150 | 153 | **ENGV** | 4 | 0.696 | 0.971 |
| 151 | 154 | **NGVT** | 4 | 0.58 | 0.986 |
| 150 | 152 | **ENG** | 3 | 1.206 | 0.834 |
| 151 | 153 | **NGV** | 3 | 0.517 | 1.011 |
| 152 | 154 | **GVT** | 3 | 0.464 | 1.055 |
| 165 | 166 | **IE** | 2 | 0.692 | 1.002 |

***Proposed epitopes.**
