## Supplemental table 2 for "Translationally controlled tumor protein *TCTP* as Peptide Vaccine against *Schistosoma japonicum*: an immunoinformatics approach"

List of epitopes that bind to MHC I:

| Peptide | Start | End | Length | Allele | ANN_ic50 | Percentile |
|---|---|---|---|---|---|---|
| **LINDVVYEV** | 22 | 30 | 9 | HLA-A*02:01 | 7.03 | 0.1 |
| | | | | HLA-A*02:06 | 14.25 | 0.2 |
| | | | | HLA-A*68:02 | 27.28 | 0.2 |
| | | | | HLA-C*15:02 | 148.97 | 4.9 |
| **RINEYMVNV** | 114 | 122 | 9 | HLA-A*02:01 | 20.73 | 0.2 |
| | | | | HLA-A*02:06 | 12.56 | 0.2 |
| | | | | HLA-A*32:01 | 238.07 | 0.2 |
| | | | | HLA-C*15:02 | 109.22 | 0.1 |
| **GLDSKLIAA** | 40 | 48 | 9 | HLA-A*02:01 | 81.49 | 0.2 |
| | | | | HLA-A*02:06 | 102.57 | 0.2 |
| **YLKGYLKAI** | 88 | 96 | 9 | HLA-A*02:01 | 144.78 | 0.2 |
| | | | | HLA-A*02:06 | 215.19 | 0.2 |
| | | | | HLA-B*08:01 | 22.97 | 0.1 |
| | | | | HLA-C*12:03 | 76.85 | 0.2 |
| | | | | HLA-C*14:02 | 144.46 | 0.2 |
| **GVTPYFVFL** | 152 | 160 | 9 | HLA-A*02:01 | 263.55 | 0.2 |
| | | | | HLA-A*02:06 | 11.7 | 0.2 |
| | | | | HLA-C*03:03 | 475.16 | 0.2 |
| **LMNFRENGV** | 145 | 153 | 9 | HLA-A*02:01 | 405.28 | 0.2 |
| **RVIDLVHAS** | 65 | 73 | 9 | HLA-A*02:06 | 156.35 | 0.2 |
| ***YMVNVFKNF** | 118 | 126 | 9 | HLA-A*02:06 | 186.59 | 0.2 |
| | | | | HLA-A*23:01 | 27.97 | 0.2 |
| | | | | HLA-A*29:02 | 229.86 | 0.2 |
| | | | | HLA-B*15:01 | 17.3 | 0.1 |
| | | | | HLA-B*15:02 | 83.03 | 0.1 |
| | | | | HLA-C*12:03 | 139.04 | 0.2 |
| | | | | HLA-C*14:02 | 56.72 | 0.2 |
| ***VVYEVDANF** | 26 | 34 | 9 | HLA-A*02:06 | 239.87 | 0.2 |
| | | | | HLA-A*23:01 | 167.66 | 0.2 |
| | | | | HLA-B*15:01 | 167.59 | 0.2 |
| | | | | HLA-B*15:02 | 478.94 | 0.1 |
| | | | | HLA-B*35:01 | 332.42 | 0.3 |
| | | | | HLA-B*58:01 | 443.26 | 0.2 |
| | | | | HLA-C*12:03 | 379.95 | 0.2 |
| | | | | HLA-C*14:02 | 365.36 | 0.2 |
| **FVFLKDGLI** | 157 | 165 | 9 | HLA-A*02:06 | 254.56 | 0.2 |
| | | | | HLA-A*68:02 | 128.17 | 0.2 |
| **FSDSHSPQL** | 14 | 22 | 9 | HLA-A*02:06 | 282.16 | 0.2 |
| | | | | HLA-B*39:01 | 258.15 | 0.2 |
| | | | | HLA-C*03:03 | 26.63 | 0.2 |

| | | | | | | |
|---|---|---|---|---|---|---|
| | | | | HLA-C*05:01 | 18.33 | 0.2 |
| | | | | HLA-C*08:02 | 148.99 | 0.1 |
| | | | | HLA-C*15:02 | 198.39 | 9.7 |
| **FLKDGLIEE** | 159 | 167 | 9 | HLA-A*02:06 | 353.46 | 0.2 |
| **RAYLKGYLK** | 86 | 94 | 9 | HLA-A*03:01 | 27.65 | 0.2 |
| | | | | HLA-A*11:01 | 23.78 | 0.2 |
| | | | | HLA-A*30:01 | 70.14 | 0.2 |
| | | | | HLA-A*31:01 | 19.97 | 0.2 |
| | | | | HLA-A*68:01 | 444.91 | 5.2 |
| **RLVSTSFDK** | 74 | 82 | 9 | HLA-A*03:01 | 90.94 | 0.2 |
| | | | | HLA-A*11:01 | 68.43 | 0.2 |
| | | | | HLA-A*30:01 | 309.08 | 0.2 |
| | | | | HLA-A*31:01 | 497.13 | 0.2 |
| **VTPYFVFLK** | 153 | 161 | 9 | HLA-A*03:01 | 102.23 | 0.2 |
| | | | | HLA-A*11:01 | 7.97 | 0.1 |
| | | | | HLA-A*30:01 | 341.09 | 0.2 |
| | | | | HLA-A*31:01 | 140.09 | 0.2 |
| | | | | HLA-A*68:01 | 12.07 | 0.1 |
| **LVSTSFDKK** | 75 | 83 | 9 | HLA-A*03:01 | 388.41 | 1.3 |
| | | | | HLA-A*11:01 | 33.2 | 0.2 |
| | | | | HLA-A*68:01 | 82.51 | 4.9 |
| **TSFDKKSYR** | 78 | 86 | 9 | HLA-A*11:01 | 70.57 | 0.2 |
| | | | | HLA-A*31:01 | 5.04 | 0.1 |
| | | | | HLA-A*68:01 | 7.16 | 0.1 |
| **KAIKERLQK** | 94 | 102 | 9 | HLA-A*11:01 | 77.86 | 0.2 |
| | | | | HLA-A*30:01 | 56.34 | 0.2 |
| **GMVALMNFR** | 141 | 149 | 9 | HLA-A*11:01 | 125.89 | 0.2 |
| | | | | HLA-A*31:01 | 10.5 | 0.1 |
| | | | | HLA-A*68:01 | 90.38 | 4.9 |
| **VYEVDANFI** | 27 | 35 | 9 | HLA-A*24:02 | 470.7 | 0.2 |
| **IFESRINEY** | 110 | 118 | 9 | HLA-A*29:02 | 448.96 | 0.2 |
| **SYRAYLKGY** | 84 | 92 | 9 | HLA-A*29:02 | 460.35 | 0.2 |
| | | | | HLA-A*30:01 | 405.43 | 0.2 |
| | | | | HLA-A*30:02 | 405.7 | 0.1 |
| | | | | HLA-B*15:02 | 382.27 | 0.1 |
| | | | | HLA-C*14:02 | 66.96 | 0.2 |
| **KSYRAYLKG** | 83 | 91 | 9 | HLA-A*30:01 | 145.03 | 0.2 |
| | | | | HLA-B*57:01 | 371.36 | 0.2 |
| | | | | HLA-B*58:01 | 341.98 | 0.2 |
| **KLIAANPSG** | 44 | 52 | 9 | HLA-A*30:01 | 226.91 | 0.2 |
| **LVHASRLVS** | 69 | 77 | 9 | HLA-A*30:01 | 248.32 | 0.2 |
| **STSFDKKSY** | 77 | 85 | 9 | HLA-A*30:02 | 77.24 | 0.1 |
| **RLQKENPER** | 99 | 107 | 9 | HLA-A*31:01 | 213.61 | 0.2 |

| | | | | | | |
|---|---|---|---|---|---|---|
| **GYLKAIKER** | 91 | 99 | 9 | HLA-A*31:01 | 374.89 | 0.2 |
| **VIDLVHASR** | 66 | 74 | 9 | HLA-A*31:01 | 411.96 | 0.2 |
| **NEYMVNVFK** | 116 | 124 | 9 | HLA-A*68:01 | 22.21 | 4.9 |
| | | | | HLA-B*18:01 | 398.43 | 0.1 |
| **ERVSIFESR** | 106 | 114 | 9 | HLA-A*68:01 | 235.42 | 5.2 |
| **EEVSDSTER** | 57 | 65 | 9 | HLA-A*68:01 | 316.48 | 5.2 |
| **EVSDSTERV** | 58 | 66 | 9 | HLA-A*68:02 | 4.44 | 0.2 |
| **ESMNPDGMV** | 135 | 143 | 9 | HLA-A*68:02 | 9.74 | 0.2 |
| | | | | HLA-C*15:02 | 331.26 | 9.7 |
| **STERVIDLV** | 62 | 70 | 9 | HLA-A*68:02 | 55.76 | 0.2 |
| **HSPQLINDV** | 18 | 26 | 9 | HLA-A*68:02 | 331.3 | 0.2 |
| **SPQLINDVV** | 19 | 27 | 9 | HLA-B*07:02 | 271.77 | 0.2 |
| **YLKAIKERL** | 92 | 100 | 9 | HLA-B*08:01 | 492.03 | 0.1 |
| **ASRLVSTSF** | 72 | 80 | 9 | HLA-B*15:01 | 44.58 | 0.1 |
| **SMNPDGMVA** | 136 | 144 | 9 | HLA-B*15:01 | 336.91 | 0.2 |
| ***YEHYIGESM** | 129 | 137 | 9 | HLA-B*15:01 | 456.89 | 0.2 |
| | | | | HLA-B*18:01 | 8.38 | 0.1 |
| | | | | HLA-B*40:01 | 9.59 | 0.1 |
| | | | | HLA-B*40:02 | 40.95 | 0.1 |
| | | | | HLA-C*03:03 | 78.58 | 0.2 |
| | | | | HLA-C*12:03 | 192.74 | 0.2 |
| | | | | HLA-C*14:02 | 49.08 | 0.2 |
| **RENGVTPYF** | 149 | 157 | 9 | HLA-B*15:01 | 475.94 | 0.2 |
| | | | | HLA-B*18:01 | 71.93 | 0.1 |
| | | | | HLA-B*40:01 | 75.05 | 0.1 |
| | | | | HLA-B*40:02 | 47.03 | 0.1 |
| | | | | HLA-B*44:02 | 245.06 | 0.1 |
| | | | | HLA-B*44:03 | 31.14 | 0.1 |
| **FDKKSYRAY** | 80 | 88 | 9 | HLA-B*15:02 | 104.99 | 0.1 |
| | | | | HLA-C*12:03 | 389.34 | 0.2 |
| ***FRENGVTPY** | 148 | 156 | 9 | HLA-B*15:02 | 131.87 | 0.1 |
| | | | | HLA-B*35:01 | 414.47 | 0.3 |
| | | | | HLA-C*06:02 | 499.61 | 0.1 |
| | | | | HLA-C*07:01 | 403.71 | 9.7 |
| | | | | HLA-C*07:02 | 68.06 | 0.1 |
| | | | | HLA-C*12:03 | 43.3 | 0.2 |
| | | | | HLA-C*14:02 | 200.98 | 0.2 |
| **FESRINEYM** | 111 | 119 | 9 | HLA-B*18:01 | 486.13 | 0.1 |
| | | | | HLA-B*40:01 | 39.74 | 0.1 |
| | | | | HLA-B*40:02 | 132 | 0.1 |
| **YRAYLKGYL** | 85 | 93 | 9 | HLA-B*27:05 | 109.53 | 0.2 |
| | | | | HLA-C*06:02 | 155.75 | 0.1 |
| | | | | HLA-C*07:01 | 168.85 | 9.7 |

| | | | | | | |
|---|---|---|---|---|---|---|
| **NPDGMVALM** | 138 | 146 | 9 | HLA-B*35:01 | 69.92 | 0.2 |
| | | | | HLA-B*39:01 | 330.36 | 0.2 |
| | | | | HLA-B*53:01 | 133.54 | 0.1 |
| **DAISGDEMF** | 6 | 14 | 9 | HLA-B*35:01 | 167.44 | 0.3 |
| **GESMNPDGM** | 134 | 142 | 9 | HLA-B*40:01 | 38.91 | 0.1 |
| **KENPERVSI** | 102 | 110 | 9 | HLA-B*40:01 | 273.26 | 0.1 |
| | | | | HLA-B*40:02 | 84.56 | 0.1 |
| | | | | HLA-B*44:02 | 479.49 | 0.1 |
| **IAANPSGEE** | 46 | 54 | 9 | HLA-C*03:03 | 229.15 | 0.2 |
| **AANPSGEEG** | 47 | 55 | 9 | HLA-C*03:03 | 301.03 | 0.2 |
| **VSDSTERVI** | 59 | 67 | 9 | HLA-C*05:01 | 118.23 | 0.2 |
| **MNPDGMVAL** | 137 | 145 | 9 | HLA-C*12:03 | 197.25 | 0.2 |
| | | | | HLA-C*14:02 | 216.16 | 0.2 |

***Proposed epitopes.**
