## Supplemental table 3 for "Translationally controlled tumor protein *TCTP* as Peptide Vaccine against *Schistosoma japonicum*: an immunoinformatics approach"

List of epitopes that bind to MHC II:

| Epitope (core sequence) | Allele | Peptide Sequence | Start | End | IC50 | Rank |
|---|---|---|---|---|---|---|
| **FDKKSYRAY** | HLA-DRB1*11:01 | STSFDKKSYRAYLKG | 77 | 91 | 64.1 | 10.23 |
| | HLA-DRB1*11:01 | TSFDKKSYRAYLKGY | 78 | 92 | 71.5 | 11.05 |
| | HLA-DRB1*11:01 | VSTSFDKKSYRAYLK | 76 | 90 | 98.2 | 13.58 |
| **FLKDGLIEE** | HLA-DPA1*02:01/DPB1*01:01 | PYFVFLKDGLIEEKY | 155 | 169 | 45.7 | 4.77 |
| | HLA-DPA1*02:01/DPB1*01:01 | TPYFVFLKDGLIEEK | 154 | 168 | 55.6 | 5.92 |
| | HLA-DPA1*02:01/DPB1*01:01 | VTPYFVFLKDGLIEE | 153 | 167 | 61.5 | 6.57 |
| | HLA-DPA1*03:01/DPB1*04:02 | PYFVFLKDGLIEEKY | 155 | 169 | 67.1 | 7.28 |
| **FRENGVTPY** | HLA-DRB1*01:01 | LMNFRENGVTPYFVF | 145 | 159 | 38.3 | 18.14 |
| | HLA-DRB1*01:01 | ALMNFRENGVTPYFV | 144 | 158 | 96.4 | 29.68 |
| | HLA-DRB3*01:01 | LMNFRENGVTPYFVF | 145 | 159 | 24.6 | 1.53 |
| | HLA-DRB3*01:01 | ALMNFRENGVTPYFV | 144 | 158 | 26.8 | 1.66 |
| | HLA-DRB3*01:01 | VALMNFRENGVTPYF | 143 | 157 | 30.5 | 1.86 |
| | HLA-DRB3*01:01 | MVALMNFRENGVTPY | 142 | 156 | 36.9 | 2.2 |
| | HLA-DRB3*01:01 | MNFRENGVTPYFVFL | 146 | 160 | 39.5 | 2.33 |
| | HLA-DRB3*01:01 | NFRENGVTPYFVFLK | 147 | 161 | 68 | 3.48 |
| **FSDSHSPQL** | HLA-DRB1*07:01 | SGDEMFSDSHSPQLI | 9 | 23 | 5.4 | 0.59 |
| | HLA-DRB1*07:01 | ISGDEMFSDSHSPQL | 8 | 22 | 5.8 | 0.68 |
| | HLA-DRB1*07:01 | GDEMFSDSHSPQLIN | 10 | 24 | 6.9 | 0.92 |
| | HLA-DRB1*07:01 | DEMFSDSHSPQLIND | 11 | 25 | 10 | 1.64 |
| | HLA-DRB1*07:01 | EMFSDSHSPQLINDV | 12 | 26 | 12.3 | 2.18 |
| | HLA-DRB1*07:01 | MFSDSHSPQLINDVV | 13 | 27 | 16.8 | 3.17 |
| | HLA-DRB1*07:01 | FSDSHSPQLINDVVY | 14 | 28 | 27.9 | 5.34 |
| | HLA-DRB1*09:01 | DEMFSDSHSPQLIND | 11 | 25 | 39.2 | 2.34 |
| | HLA-DRB1*09:01 | GDEMFSDSHSPQLIN | 10 | 24 | 47.6 | 3.04 |
| | HLA-DRB1*09:01 | EMFSDSHSPQLINDV | 12 | 26 | 51.5 | 3.35 |
| | HLA-DRB1*09:01 | SGDEMFSDSHSPQLI | 9 | 23 | 64.8 | 4.37 |
| | HLA-DRB1*09:01 | MFSDSHSPQLINDVV | 13 | 27 | 84.8 | 5.87 |
| | HLA-DRB1*09:01 | ISGDEMFSDSHSPQL | 8 | 22 | 98.7 | 6.82 |
| ***FVFLKDGLI** | HLA-DRB1*01:01 | TPYFVFLKDGLIEEK | 154 | 168 | 4.7 | 0.54 |
| | HLA-DRB1*01:01 | PYFVFLKDGLIEEKY | 155 | 169 | 5.3 | 1.06 |
| | HLA-DRB1*01:01 | VTPYFVFLKDGLIEE | 153 | 167 | 5.5 | 1.24 |
| | HLA-DRB1*01:01 | GVTPYFVFLKDGLIE | 152 | 166 | 6 | 1.71 |
| | HLA-DRB1*01:01 | NGVTPYFVFLKDGLI | 151 | 165 | 6.7 | 2.37 |
| | HLA-DRB1*04:05 | VTPYFVFLKDGLIEE | 153 | 167 | 49.2 | 4.77 |
| | HLA-DRB1*04:05 | TPYFVFLKDGLIEEK | 154 | 168 | 52.7 | 5.15 |
| | HLA-DRB1*04:05 | GVTPYFVFLKDGLIE | 152 | 166 | 55.2 | 5.41 |
| | HLA-DRB1*04:05 | NGVTPYFVFLKDGLI | 151 | 165 | 61.5 | 6.06 |
| | HLA-DRB1*04:05 | PYFVFLKDGLIEEKY | 155 | 169 | 68.8 | 6.78 |
| | HLA-DRB1*07:01 | NGVTPYFVFLKDGLI | 151 | 165 | 11 | 1.88 |

| | | | | | | |
|---|---|---|---|---|---|---|
| | HLA-DRB1*07:01 | GVTPYFVFLKDGLIE | 152 | 166 | 16.4 | 3.08 |
| | HLA-DRB1*07:01 | VTPYFVFLKDGLIEE | 153 | 167 | 26.3 | 5.07 |
| | HLA-DRB1*07:01 | TPYFVFLKDGLIEEK | 154 | 168 | 43.1 | 7.69 |
| | HLA-DRB1*07:01 | PYFVFLKDGLIEEKY | 155 | 169 | 61.5 | 10.12 |
| | HLA-DRB5*01:01 | TPYFVFLKDGLIEEK | 154 | 168 | 13.5 | 3.32 |
| | HLA-DRB5*01:01 | VTPYFVFLKDGLIEE | 153 | 167 | 17.3 | 4.26 |
| | HLA-DRB5*01:01 | PYFVFLKDGLIEEKY | 155 | 169 | 17.8 | 4.38 |
| | HLA-DRB5*01:01 | GVTPYFVFLKDGLIE | 152 | 166 | 21 | 5.1 |
| | HLA-DRB5*01:01 | NGVTPYFVFLKDGLI | 151 | 165 | 25.8 | 6.1 |
| ***IDLVHASRL** | HLA-DRB1*01:01 | ERVIDLVHASRLVST | 64 | 78 | 14.7 | 8.42 |
| | HLA-DRB1*01:01 | TERVIDLVHASRLVS | 63 | 77 | 18.3 | 10.44 |
| | HLA-DRB1*01:01 | RVIDLVHASRLVSTS | 65 | 79 | 18.4 | 10.49 |
| | HLA-DRB1*01:01 | STERVIDLVHASRLV | 62 | 76 | 23.3 | 12.82 |
| | HLA-DRB1*01:01 | DSTERVIDLVHASRL | 61 | 75 | 32.3 | 16.25 |
| | HLA-DRB1*01:01 | VIDLVHASRLVSTSF | 66 | 80 | 33 | 16.49 |
| | HLA-DRB1*01:01 | IDLVHASRLVSTSFD | 67 | 81 | 47.3 | 20.58 |
| | HLA-DRB1*03:01 | ERVIDLVHASRLVST | 64 | 78 | 68.1 | 3.71 |
| | HLA-DRB1*03:01 | TERVIDLVHASRLVS | 63 | 77 | 70.3 | 3.8 |
| | HLA-DRB1*03:01 | STERVIDLVHASRLV | 62 | 76 | 87.6 | 4.51 |
| | HLA-DRB1*07:01 | STERVIDLVHASRLV | 62 | 76 | 3 | 0.1 |
| | HLA-DRB1*07:01 | TERVIDLVHASRLVS | 63 | 77 | 3.1 | 0.11 |
| | HLA-DRB1*07:01 | ERVIDLVHASRLVST | 64 | 78 | 3.3 | 0.15 |
| | HLA-DRB1*07:01 | DSTERVIDLVHASRL | 61 | 75 | 3.6 | 0.2 |
| | HLA-DRB1*07:01 | RVIDLVHASRLVSTS | 65 | 79 | 3.7 | 0.21 |
| | HLA-DRB1*07:01 | VIDLVHASRLVSTSF | 66 | 80 | 4.9 | 0.48 |
| | HLA-DRB1*07:01 | IDLVHASRLVSTSFD | 67 | 81 | 6.5 | 0.83 |
| | HLA-DRB1*09:01 | ERVIDLVHASRLVST | 64 | 78 | 82.7 | 5.72 |
| | HLA-DRB1*13:02 | TERVIDLVHASRLVS | 63 | 77 | 27.4 | 1.99 |
| | HLA-DRB1*13:02 | STERVIDLVHASRLV | 62 | 76 | 28 | 2.03 |
| | HLA-DRB1*13:02 | ERVIDLVHASRLVST | 64 | 78 | 28.7 | 2.08 |
| | HLA-DRB1*13:02 | DSTERVIDLVHASRL | 61 | 75 | 37.6 | 2.69 |
| | HLA-DRB1*13:02 | RVIDLVHASRLVSTS | 65 | 79 | 48.4 | 3.38 |
| | HLA-DRB1*13:02 | VIDLVHASRLVSTSF | 66 | 80 | 80.8 | 5.2 |
| | HLA-DRB1*15:01 | ERVIDLVHASRLVST | 64 | 78 | 41.1 | 4.05 |
| | HLA-DRB1*15:01 | TERVIDLVHASRLVS | 63 | 77 | 42.3 | 4.19 |
| | HLA-DRB1*15:01 | STERVIDLVHASRLV | 62 | 76 | 50.7 | 5.17 |
| | HLA-DRB1*15:01 | RVIDLVHASRLVSTS | 65 | 79 | 51.2 | 5.22 |
| | HLA-DRB1*15:01 | DSTERVIDLVHASRL | 61 | 75 | 78.9 | 8.04 |
| | HLA-DRB1*15:01 | VIDLVHASRLVSTSF | 66 | 80 | 84.5 | 8.56 |
| | HLA-DRB5*01:01 | TERVIDLVHASRLVS | 63 | 77 | 59.4 | 11.18 |
| | HLA-DRB5*01:01 | STERVIDLVHASRLV | 62 | 76 | 69.8 | 12.38 |
| | HLA-DRB5*01:01 | ERVIDLVHASRLVST | 64 | 78 | 74.9 | 12.94 |
| **INEYMVNVF** | HLA-DPA1*02:01/DPB1*01:01 | ESRINEYMVNVFKNF | 112 | 126 | 50.4 | 5.32 |

| | | | | | | |
|---|---|---|---|---|---|---|
| **KKSYRAYLK** | HLA-DPA1*02:01/DPB1*05:01 | FDKKSYRAYLKGYLK | 80 | 94 | 72 | 1.29 |
| | HLA-DRB5*01:01 | SFDKKSYRAYLKGYL | 79 | 93 | 11.5 | 2.79 |
| | HLA-DRB5*01:01 | TSFDKKSYRAYLKGY | 78 | 92 | 19 | 4.64 |
| | HLA-DRB5*01:01 | STSFDKKSYRAYLKG | 77 | 91 | 26.2 | 6.17 |
| | HLA-DRB5*01:01 | VSTSFDKKSYRAYLK | 76 | 90 | 28.7 | 6.63 |
| **LINDVVYEV** | HLA-DRB1*03:01 | SPQLINDVVYEVDAN | 19 | 33 | 45.4 | 2.65 |
| | HLA-DRB1*03:01 | HSPQLINDVVYEVDA | 18 | 32 | 51.6 | 2.95 |
| | HLA-DRB1*03:01 | SHSPQLINDVVYEVD | 17 | 31 | 72.1 | 3.87 |
| | HLA-DRB1*03:01 | PQLINDVVYEVDANF | 20 | 34 | 88.3 | 4.54 |
| | HLA-DRB3*01:01 | SHSPQLINDVVYEVD | 17 | 31 | 23.6 | 1.47 |
| | HLA-DRB3*01:01 | DSHSPQLINDVVYEV | 16 | 30 | 25.2 | 1.57 |
| | HLA-DRB3*01:01 | HSPQLINDVVYEVDA | 18 | 32 | 25.3 | 1.57 |
| | HLA-DRB3*01:01 | SPQLINDVVYEVDAN | 19 | 33 | 28.4 | 1.74 |
| | HLA-DRB3*01:01 | PQLINDVVYEVDANF | 20 | 34 | 57.1 | 3.07 |
| **LKAIKERLQ** | HLA-DRB1*01:01 | KGYLKAIKERLQKEN | 90 | 104 | 52.5 | 21.84 |
| **LKGYLKAIK** | HLA-DRB1*04:05 | RAYLKGYLKAIKERL | 86 | 100 | 72.9 | 7.17 |
| | HLA-DRB1*15:01 | RAYLKGYLKAIKERL | 86 | 100 | 52.7 | 5.38 |
| | HLA-DRB1*15:01 | AYLKGYLKAIKERLQ | 87 | 101 | 61.2 | 6.3 |
| | HLA-DRB5*01:01 | RAYLKGYLKAIKERL | 86 | 100 | 4.3 | 0.63 |
| **LQKENPERV** | HLA-DRB1*13:02 | KERLQKENPERVSIF | 97 | 111 | 88.2 | 5.55 |
| **LVHASRLVS** | HLA-DRB1*03:01 | VIDLVHASRLVSTSF | 66 | 80 | 80.7 | 4.23 |
| | HLA-DRB1*03:01 | RVIDLVHASRLVSTS | 65 | 79 | 94.4 | 4.82 |
| | HLA-DRB1*11:01 | VIDLVHASRLVSTSF | 66 | 80 | 7.6 | 0.76 |
| | HLA-DRB1*11:01 | RVIDLVHASRLVSTS | 65 | 79 | 10.3 | 1.4 |
| | HLA-DRB1*11:01 | IDLVHASRLVSTSFD | 67 | 81 | 10.3 | 1.4 |
| | HLA-DRB1*11:01 | ERVIDLVHASRLVST | 64 | 78 | 15.3 | 2.55 |
| | HLA-DRB1*11:01 | DLVHASRLVSTSFDK | 68 | 82 | 15.6 | 2.62 |
| | HLA-DRB1*11:01 | TERVIDLVHASRLVS | 63 | 77 | 24.2 | 4.44 |
| | HLA-DRB1*11:01 | LVHASRLVSTSFDKK | 69 | 83 | 25.7 | 4.73 |
| **LVSTSFDKK** | HLA-DPA1*01:03/DPB1*02:01 | HASRLVSTSFDKKSY | 71 | 85 | 30.7 | 3.5 |
| | HLA-DPA1*01:03/DPB1*02:01 | VHASRLVSTSFDKKS | 70 | 84 | 32.3 | 3.65 |
| | HLA-DPA1*01:03/DPB1*02:01 | LVHASRLVSTSFDKK | 69 | 83 | 33.9 | 3.79 |
| | HLA-DPA1*01:03/DPB1*02:01 | ASRLVSTSFDKKSYR | 72 | 86 | 35.3 | 3.92 |
| | HLA-DPA1*01:03/DPB1*02:01 | SRLVSTSFDKKSYRA | 73 | 87 | 67.8 | 6.4 |
| **MFSDSHSPQ** | HLA-DRB1*04:01 | GDEMFSDSHSPQLIN | 10 | 24 | 65 | 5.19 |
| | HLA-DRB1*04:01 | DEMFSDSHSPQLIND | 11 | 25 | 75.3 | 6.08 |
| | HLA-DRB1*04:01 | SGDEMFSDSHSPQLI | 9 | 23 | 97.7 | 7.93 |
| | HLA-DRB1*04:01 | EMFSDSHSPQLINDV | 12 | 26 | 99.7 | 8.08 |
| **MVALMNFRE** | HLA-DRB1*04:04 | PDGMVALMNFRENGV | 139 | 153 | 54.6 | 6.32 |
| | HLA-DRB1*04:04 | DGMVALMNFRENGVT | 140 | 154 | 57.5 | 6.71 |
| | HLA-DRB1*04:04 | GMVALMNFRENGVTP | 141 | 155 | 61.2 | 7.25 |
| | HLA-DRB1*04:04 | NPDGMVALMNFRENG | 138 | 152 | 61.5 | 7.28 |
| | HLA-DRB1*04:04 | MNPDGMVALMNFREN | 137 | 151 | 82.9 | 9.98 |

| | | | | | | |
|---|---|---|---|---|---|---|
| | HLA-DRB1*04:05 | PDGMVALMNFRENGV | 139 | 153 | 81.5 | 8 |
| | HLA-DRB1*15:01 | PDGMVALMNFRENGV | 139 | 153 | 72.6 | 7.44 |
| | HLA-DRB1*15:01 | DGMVALMNFRENGVT | 140 | 154 | 80.5 | 8.19 |
| | HLA-DRB1*15:01 | MNPDGMVALMNFREN | 137 | 151 | 91.3 | 9.17 |
| | HLA-DRB1*15:01 | GMVALMNFRENGVTP | 141 | 155 | 96.4 | 9.62 |
| **PDGMVALMN** | HLA-DQA1*05:01/DQB1*03:01 | SMNPDGMVALMNFRE | 136 | 150 | 80.9 | 12.87 |
| **PDGMVALMN** | HLA-DQA1*05:01/DQB1*03:01 | ESMNPDGMVALMNFR | 135 | 149 | 84.2 | 13.23 |
| **RAYLKGYLK** | HLA-DRB5*01:01 | KSYRAYLKGYLKAIK | 83 | 97 | 3.6 | 0.42 |
| | HLA-DRB5*01:01 | KKSYRAYLKGYLKAI | 82 | 96 | 3.9 | 0.51 |
| | HLA-DRB5*01:01 | DKKSYRAYLKGYLKA | 81 | 95 | 4.6 | 0.72 |
| | HLA-DRB5*01:01 | SYRAYLKGYLKAIKE | 84 | 98 | 4.7 | 0.75 |
| | HLA-DRB5*01:01 | FDKKSYRAYLKGYLK | 80 | 94 | 5.2 | 0.91 |
| | HLA-DRB5*01:01 | YRAYLKGYLKAIKER | 85 | 99 | 5.3 | 0.93 |
| **RLVSTSFDK** | HLA-DPA1*01/DPB1*04:01 | ASRLVSTSFDKKSYR | 72 | 86 | 64.5 | 3.78 |
| | HLA-DPA1*01/DPB1*04:01 | HASRLVSTSFDKKSY | 71 | 85 | 65.3 | 3.82 |
| | HLA-DPA1*01/DPB1*04:01 | VHASRLVSTSFDKKS | 70 | 84 | 75.9 | 4.3 |
| **SKLIAANPS** | HLA-DRB1*01:01 | GLDSKLIAANPSGEE | 40 | 54 | 46.3 | 20.33 |
| | HLA-DRB1*01:01 | NGLDSKLIAANPSGE | 39 | 53 | 62 | 23.9 |
| | HLA-DRB1*01:01 | SNGLDSKLIAANPSG | 38 | 52 | 85.5 | 28.05 |
| | HLA-DRB1*01:01 | LDSKLIAANPSGEEG | 41 | 55 | 87.1 | 28.3 |
| **TPYFVFLKD** | HLA-DRB1*04:04 | VTPYFVFLKDGLIEE | 153 | 167 | 95.8 | 11.43 |
| | HLA-DRB1*04:04 | TPYFVFLKDGLIEEK | 154 | 168 | 98.3 | 11.69 |
| **VHASRLVST** | HLA-DQA1*01:02/DQB1*06:02 | IDLVHASRLVSTSFD | 67 | 81 | 26.8 | 1.03 |
| | HLA-DQA1*01:02/DQB1*06:02 | VIDLVHASRLVSTSF | 66 | 80 | 29.2 | 1.21 |
| | HLA-DQA1*01:02/DQB1*06:02 | DLVHASRLVSTSFDK | 68 | 82 | 36.4 | 1.79 |
| | HLA-DQA1*01:02/DQB1*06:02 | RVIDLVHASRLVSTS | 65 | 79 | 45.2 | 2.53 |
| | HLA-DQA1*01:02/DQB1*06:02 | LVHASRLVSTSFDKK | 69 | 83 | 57.3 | 3.57 |
| | HLA-DQA1*05:01/DQB1*03:01 | VIDLVHASRLVSTSF | 66 | 80 | 14.2 | 2.42 |
| | HLA-DQA1*05:01/DQB1*03:01 | IDLVHASRLVSTSFD | 67 | 81 | 14.2 | 2.42 |
| | HLA-DQA1*05:01/DQB1*03:01 | RVIDLVHASRLVSTS | 65 | 79 | 15.3 | 2.66 |
| | HLA-DQA1*05:01/DQB1*03:01 | ERVIDLVHASRLVST | 64 | 78 | 21.2 | 3.98 |
| | HLA-DQA1*05:01/DQB1*03:01 | DLVHASRLVSTSFDK | 68 | 82 | 22.7 | 4.29 |
| | HLA-DQA1*05:01/DQB1*03:01 | LVHASRLVSTSFDKK | 69 | 83 | 59.7 | 10.31 |
| | HLA-DQA1*05:01/DQB1*03:01 | VHASRLVSTSFDKKS | 70 | 84 | 68.3 | 11.4 |
| **VVYEVDANF** | HLA-DQA1*05:01/DQB1*02:01 | QLINDVVYEVDANFI | 21 | 35 | 38.1 | 0.32 |
| | HLA-DQA1*05:01/DQB1*02:01 | PQLINDVVYEVDANF | 20 | 34 | 55.5 | 0.68 |
| **VYEVDANFI** | HLA-DRB1*07:01 | QLINDVVYEVDANFI | 21 | 35 | 26.5 | 5.09 |
| | HLA-DRB1*13:02 | QLINDVVYEVDANFI | 21 | 35 | 75.1 | 4.91 |
| **YFVFLKDGL** | HLA-DPA1*01/DPB1*04:01 | VTPYFVFLKDGLIEE | 153 | 167 | 85.4 | 4.72 |
| | HLA-DRB1*11:01 | TPYFVFLKDGLIEEK | 154 | 168 | 72.8 | 11.19 |
| | HLA-DRB1*11:01 | VTPYFVFLKDGLIEE | 153 | 167 | 89.2 | 12.79 |
| | HLA-DRB1*11:01 | PYFVFLKDGLIEEKY | 155 | 169 | 97.9 | 13.55 |
| **YIGESMNPD** | HLA-DRB1*04:05 | YEHYIGESMNPDGMV | 129 | 143 | 30.6 | 2.55 |

| | | | | | | |
|---|---|---|---|---|---|---|
| | HLA-DRB1*04:05 | DYEHYIGESMNPDGM | 128 | 142 | 34.2 | 3.01 |
| | HLA-DRB1*04:05 | EHYIGESMNPDGMVA | 130 | 144 | 47.9 | 4.62 |
| | HLA-DRB1*04:05 | HYIGESMNPDGMVAL | 131 | 145 | 70.7 | 6.96 |
| ***YLKAIKERL** | HLA-DRB1*01:01 | LKGYLKAIKERLQKE | 89 | 103 | 34.6 | 17.01 |
| | HLA-DRB1*01:01 | YLKGYLKAIKERLQK | 88 | 102 | 39.8 | 18.58 |
| | HLA-DRB1*04:05 | AYLKGYLKAIKERLQ | 87 | 101 | 84.7 | 8.31 |
| | HLA-DRB1*04:05 | YLKGYLKAIKERLQK | 88 | 102 | 96.3 | 9.31 |
| | HLA-DRB1*07:01 | RAYLKGYLKAIKERL | 86 | 100 | 66.8 | 10.76 |
| | HLA-DRB1*07:01 | AYLKGYLKAIKERLQ | 87 | 101 | 72.3 | 11.4 |
| | HLA-DRB1*07:01 | YLKGYLKAIKERLQK | 88 | 102 | 97.6 | 13.96 |
| | HLA-DRB1*09:01 | LKGYLKAIKERLQKE | 89 | 103 | 69.5 | 4.72 |
| | HLA-DRB1*09:01 | RAYLKGYLKAIKERL | 86 | 100 | 79.9 | 5.5 |
| | HLA-DRB1*09:01 | AYLKGYLKAIKERLQ | 87 | 101 | 93.2 | 6.46 |
| | HLA-DRB1*09:01 | KGYLKAIKERLQKEN | 90 | 104 | 93.5 | 6.48 |
| | HLA-DRB1*09:01 | YLKGYLKAIKERLQK | 88 | 102 | 96.2 | 6.65 |
| | HLA-DRB1*11:01 | LKGYLKAIKERLQKE | 89 | 103 | 9.1 | 1.11 |
| | HLA-DRB1*11:01 | YLKGYLKAIKERLQK | 88 | 102 | 11.5 | 1.68 |
| | HLA-DRB1*11:01 | KGYLKAIKERLQKEN | 90 | 104 | 11.5 | 1.68 |
| | HLA-DRB1*11:01 | AYLKGYLKAIKERLQ | 87 | 101 | 17.2 | 2.96 |
| | HLA-DRB1*11:01 | GYLKAIKERLQKENP | 91 | 105 | 17.9 | 3.13 |
| | HLA-DRB1*11:01 | RAYLKGYLKAIKERL | 86 | 100 | 23 | 4.2 |
| | HLA-DRB1*11:01 | YLKAIKERLQKENPE | 92 | 106 | 32 | 5.86 |
| | HLA-DRB1*15:01 | YLKGYLKAIKERLQK | 88 | 102 | 59.7 | 6.14 |
| | HLA-DRB1*15:01 | LKGYLKAIKERLQKE | 89 | 103 | 88.3 | 8.9 |
| | HLA-DRB5*01:01 | YLKGYLKAIKERLQK | 88 | 102 | 5.3 | 0.93 |
| | HLA-DRB5*01:01 | LKGYLKAIKERLQKE | 89 | 103 | 5.4 | 0.97 |
| | HLA-DRB5*01:01 | AYLKGYLKAIKERLQ | 87 | 101 | 5.7 | 1.07 |
| | HLA-DRB5*01:01 | KGYLKAIKERLQKEN | 90 | 104 | 8.3 | 1.86 |
| | HLA-DRB5*01:01 | GYLKAIKERLQKENP | 91 | 105 | 14 | 3.46 |
| | HLA-DRB5*01:01 | YLKAIKERLQKENPE | 92 | 106 | 24.5 | 5.84 |
| **YLKGYLKAI** | HLA-DRB1*01:01 | YRAYLKGYLKAIKER | 85 | 99 | 11.4 | 6.24 |
| | HLA-DRB1*01:01 | SYRAYLKGYLKAIKE | 84 | 98 | 18.3 | 10.44 |
| | HLA-DRB1*01:01 | RAYLKGYLKAIKERL | 86 | 100 | 20.8 | 11.69 |
| | HLA-DRB1*01:01 | KSYRAYLKGYLKAIK | 83 | 97 | 20.9 | 11.74 |
| | HLA-DRB1*01:01 | KKSYRAYLKGYLKAI | 82 | 96 | 34.8 | 17.08 |
| | HLA-DRB1*01:01 | AYLKGYLKAIKERLQ | 87 | 101 | 42.4 | 19.3 |
| **YMVNVFKNF** | HLA-DPA1*01/DPB1*04:01 | ESRINEYMVNVFKNF | 112 | 126 | 64.9 | 3.81 |
| | HLA-DPA1*01:03/DPB1*02:01 | ESRINEYMVNVFKNF | 112 | 126 | 24.5 | 2.87 |
| **YRAYLKGYL** | HLA-DPA1*02:01/DPB1*05:01 | KKSYRAYLKGYLKAI | 82 | 96 | 47.5 | 0.7 |
| | HLA-DPA1*02:01/DPB1*05:01 | DKKSYRAYLKGYLKA | 81 | 95 | 57.8 | 0.95 |
| | HLA-DPA1*02:01/DPB1*05:01 | KSYRAYLKGYLKAIK | 83 | 97 | 60.5 | 1.01 |
| | HLA-DPA1*02:01/DPB1*05:01 | SYRAYLKGYLKAIKE | 84 | 98 | 69.1 | 1.22 |
| | HLA-DRB1*11:01 | KKSYRAYLKGYLKAI | 82 | 96 | 15 | 2.48 |

| | | | | | | |
|---|---|---|---|---|---|---|
| | HLA-DRB1*11:01 | KSYRAYLKGYLKAIK | 83 | 97 | 19.2 | 3.41 |
| | HLA-DRB1*11:01 | DKKSYRAYLKGYLKA | 81 | 95 | 23.8 | 4.36 |
| | HLA-DRB1*11:01 | SYRAYLKGYLKAIKE | 84 | 98 | 30.2 | 5.55 |
| | HLA-DRB1*11:01 | YRAYLKGYLKAIKER | 85 | 99 | 35.2 | 6.38 |
| | HLA-DRB1*11:01 | FDKKSYRAYLKGYLK | 80 | 94 | 37.1 | 6.67 |
| | HLA-DRB1*11:01 | SFDKKSYRAYLKGYL | 79 | 93 | 53.8 | 8.99 |
| | HLA-DRB1*15:01 | KKSYRAYLKGYLKAI | 82 | 96 | 10.6 | 0.47 |
| | HLA-DRB1*15:01 | DKKSYRAYLKGYLKA | 81 | 95 | 12.2 | 0.63 |
| | HLA-DRB1*15:01 | KSYRAYLKGYLKAIK | 83 | 97 | 13.4 | 0.75 |
| | HLA-DRB1*15:01 | FDKKSYRAYLKGYLK | 80 | 94 | 13.7 | 0.78 |
| | HLA-DRB1*15:01 | SFDKKSYRAYLKGYL | 79 | 93 | 16.2 | 1.06 |
| | HLA-DRB1*15:01 | SYRAYLKGYLKAIKE | 84 | 98 | 16.9 | 1.15 |
| | HLA-DRB1*15:01 | YRAYLKGYLKAIKER | 85 | 99 | 21.9 | 1.76 |

***Proposed epitopes.**
